# The fork restart factor PHF6 interacts with RRM2 and binds to H3K56ac marked nascent DNA

**DOI:** 10.1101/2023.03.08.531704

**Authors:** Lisa Depestel, Sarah-Lee Bekaert, Ellen Sanders, Carolien Van Damme, Aline Eggermont, Siebe Loontiens, Laurentijn Tilleman, Filip Van Nieuwerburgh, Louis Delhaye, Pieter Van Vlierberghe, Sven Eyckerman, Frank Speleman, Kaat Durinck

## Abstract

The PHF6 protein is a presumed chromatin reader implicated in disease through germline loss-of-function mutations causing cognitive disability syndromes and somatic mutations are predominantly observed in acute T-cell leukemia. Previous reports support a role for PHF6 in DNA damage repair, replication fork restart as well as hematopoietic precursor cell self-renewal capacity and lineage commitment. To explore better how PHF6 mediates these functions, we mapped the PHF6 interactome and identified RRM2 as a consistent binding partner across different normal and malignant cell types. Next, PHF6 knockdown imposed increased replicative stress/DNA damage and suggested possible binding of PHF6 to H3K56ac, a marker for nascent DNA at sites of DNA damage repair. Genome-wide mapping of PHF6 chromatin binding indeed revealed overlap with sites of active DNA damage, binding sites of replication fork proteins and functional crosstalk with the neuroblastoma transcription core regulatory circuitry. Altogether, we show a canonical PHF6-RRM2 interaction enabling active transport of RRM2 to genomic sites of PHF6 mediated fork restart and PHF6 localization to H3K56ac at highly transcribed genes facilitating fork restart following replication-transcription conflicts.

## Introduction

Previous studies have shown a functional connection between replicative stress resistance, DNA damage repair and nucleosome remodeling in control of chromatin dynamics (Kotsantis *et al*, 2018; Rother & van Attikum, 2017). The chromatin reader ‘plant homeodomain zinc finger protein 6’ (PHF6) is an interactor of various well-established nucleosome remodeling complexes including NuRD (Todd & Picketts, 2012), PAF1 (Zhang *et al*, 2013) and SWI-SNF (Alvarez *et al*, 2022) and has been shown to mediate replication fork restart and DNA damage repair (Alvarez *et al*, 2022; Warmerdam *et al*, 2019). However, the histone post-translational modifications (PTMs) recognized and bound by PHF6 as well as the molecular basis for the functional involvement of PHF6 in these processes still remain elusive. The PHF6 protein structure contains four nuclear/nucleolar localization signals (NLS) as well as two nearly identical extended PHD-like zinc finger domains (ePHD). In addition to PTMs, these ePHD domains also mediate protein-protein interactions (Aasland et al., 1995; Mellor, 2006).

Mutations in *PHF6* were first described in the Börjeson-Forssman-Lehman syndrome (BFLS), an X-linked mental retardation disorder (Chao *et al*, 2010; Lower *et al*, 2002). Later, *PHF6* mutations were also identified in patients with the Coffin-Siris syndrome (Wieczorek *et al*, 2013), which show overlapping phenotypic features with BFLS and in which mainly mutations in several components of the SWI/SNF chromatin remodeling complex were previously reported. These observations suggest a critical role for PHF6 in neuronal development, which is further supported by the high expression of *Phf6* in the developing central nervous system in mice and its functions in regulation of neuronal proliferation, neurite outgrowth and migration by interacting with the PAF1 complex (Zhang *et al*, 2013; Fliedner *et al*, 2020; Franzoni *et al*, 2015; Voss *et al*, 2007). Of further interest, previous work pointed to a key role for PHF6 as a tumor suppressor, predominantly in T-cell acute lymphoblastic leukemia (T-ALL) (Van Vlierberghe *et al*, 2010). Later, mutations were also reported in other cancer entities including AML (Van Vlierberghe *et al*, 2011), CML (Li *et al*, 2013), mixed phenotype acute leukemia (MPAL) (Xiao *et al*, 2018) and hepatocellular carcinoma (Yoo et al., 2012). In contrast to this apparent tumor suppressor role, PHF6 is overexpressed in other cancer types including breast cancer, colorectal cancers and T-ALL without evidence for PHF6 mutations (Hajjari *et al*, 2016), suggesting that PHF6 may act both as either tumor suppressor or tumor promoting gene depending on the cellular lineage context.

In order to understand how PHF6 is linked to these seemingly unrelated functionalities, we followed an unbiased approach by mapping the PHF6 interactome in normal and malignant cells. Surprisingly, RRM2 emerged as a consistent interaction partner. RRM2 forms heterodimers with RRM1 and acts as catalytic subunit of the ribonucleotide reductase (RNR) enzyme, which produces essential dNTPs for DNA replication/repair and is critically regulated under replicative stress by the ATR-CHK1 pathway. Using a bioluminescence resonance energy transfer (BRET)-based assay, we confirmed the PHF6-RRM2 complex and mapped the interaction interface. Using neuroblastoma cells with high PHF6 and RRM2 expression levels, we revealed overlap of PHF6 genome-wide binding sites with DNA damage sites and chromatin co-localization with several proteins acting at replication forks. Transcriptome profiling following PHF6 knockdown suggested a functional connection of PHF6 with H3K56ac, a known nascent DNA chromatin mark which suppresses break-induced DNA repair and controls expression homeostasis during S-phase (Whale *et al*, 2022). We confirmed the potential of PHF6 to bind at this epigenetic mark and high density of H3K56ac at the coding region of highly transcribed neuroblastoma core regulatory circuit lineage specific transcription factors.

## Results

### PHF6 interacts with RRM2 in multiple cancer and non-malignant cell lines

In follow-up of our previous work on PHF6 in the context of T-ALL (Van Vlierberghe *et al*, 2010; Loontiens *et al*, 2020), we aimed to gain further insight into the functional role of PHF6 through identification of its interacting partners. For this interactome analysis, we applied immunoprecipitation coupled mass spectrometry (IP-MS) in HEK293T cells using endogenous PHF6 as bait and identified the ribonucleotide reductase subunit M2 (RRM2) as top-ranked PHF6 interaction partner (**Figure 1A**). RRM2 is part of the ribonucleotide reductase complex (RNR), the rate-limiting enzyme involved in the synthesis of dNTPs and involved in replication, DNA damage and replicative stress responses (Nunes *et al*, 2022). Given the hitherto unexplained role of PHF6 in fork restart (Warmerdam *et al*, 2019), we hypothesize that the PHF6-RRM2 interaction could be crucial to provide sufficient local dNTPs, thereby facilitating timely fork restart at genomic regions affected by replicative stress. In view of our recent discovery of RRM2 as an important replicative stress resistance dependency factor in high-risk neuroblastoma (Nunes *et al*, 2022) and high PHF6 expression levels in this tumor entity, we decided to further investigate the role of PHF6 and PHF6-RRM2 interaction in neuroblastoma cell lines. IP-MS experiments in *MYCN* amplified (IMR-32, SK-N-BE(2)-C)) and *MYCN* non-amplified (CLB-GA, SH-SY5Y) neuroblastoma cell lines, confirmed consistent interaction of PHF6 with RRM2 (**Figure 1B and S1**). Furthermore, we further validated this interaction in additional tumor entities and cellular contexts including acute T-cell leukemia (ALL-SIL), colorectal cancer (HCT116) as well as normal immortalized retinal pigmented epithelial cells (RPE) (**Figure S1**).

**Figure 1.**
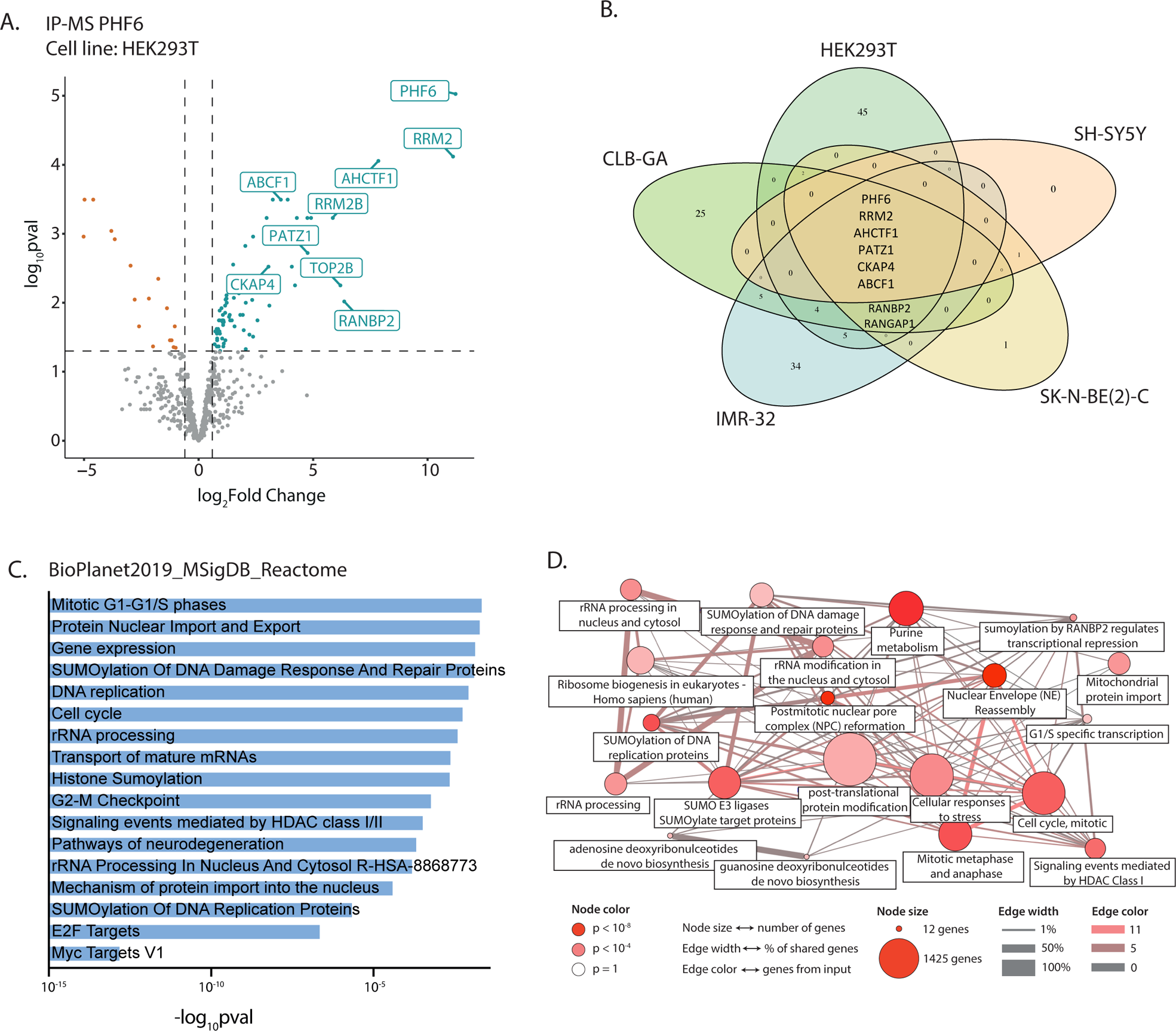
PHF6 interacts with RRM2 and proteins involved in nuclear transport, SUMOylation and cell cycle regulation. **(A)** Volcano plot displaying significantly enriched putative PHF6 interactors by means of immunoprecipitation-coupled mass spectrometry (IP-MS) in HEK293T cells; **(B)** Venn diagram showing PHF6 interactors commonly identified between IP-MS experiments in HEK293T and CLB-GA, IMR-32, SK-N-BE(2)-C and SH-SY5Y neuroblastoma cell lines; **(C)** ENRICHR and **(D)** ConsensusPathDB over-representation analysis showing functional enrichment of PHF6 interacting proteins in processes such as cell cycle regulation, gene expression, rRNA processing, SUMOylation, nuclear transport and replication.

Gene Ontology (GO) analysis for the PHF6 interactor hit list (FDR < 0.05) using ENRICHR (Xie *et al*, 2021) and ConsensusPathDB (Kamburov *et al*, 2009) tools showed for proteins involved in ribosomal RNA processing, cell cycle regulation, DNA repair and HDAC1/2 mediated signaling events, in accordance with previously reported functions of PHF6 (Todd et al., 2015). Interestingly, also proteins involved in nuclear transport of proteins and maturation of mRNA, as well as SUMOylation, a PTM associated with nuclear protein import (Hay, 2005), were enriched (**Figure 1C and 1D**). Taken together, we identify RRM2 as a consistent PHF6 interacting protein across multiple cell types suggesting a critical role for PHF6 bound RRM2 in PHF6 mediated DNA repair processes.

### PHF6 binds to RRM2 through its first PHD domain

Endogenous co-immunoprecipitation (co-IP) using PHF6 as bait, followed by immunoblotting for RRM2 and PHF6 in HEK293T and IMR-32 neuroblastoma cells, confirmed the protein-protein interaction of PHF6 with RRM2 as observed by IP-MS (**Figure 2A**). Next, we used the bioluminescence resonance energy transfer (NanoBRET™) technology (Dale *et al*, 2019) as orthogonal method to further investigate the PHF6-RRM2 interaction (**Figure 2B**). In brief, plasmids with N-terminal fusion of the NanoLuc® luciferase to PHF6 (Nluc-PHF6) and a C-terminal fusion of the HaloTag® to RRM2 (RRM2-HT) were co-transfected in HEK293T cells and resulted in the highest BRET ratios (**Figure S2A**), which were not significantly dependent on the amount of transfected plasmids (**Figure 2C**). The donor saturation assay curve of Nluc-PHF6 and RRM2-HT reaches hyperbolic saturation (**Figure 2D**), confirming the specificity of the interaction. The protein structure of PHF6 is characterized by two zinc finger domains called ‘<u>z</u>inc knuckle, <u>a</u>typical <u>P</u>HD’ (ZaP) domains and four localization signals for the nucleus and nucleoli (N(o)LS). Since this atypical PHD domain is unstable in solution, the N-terminal pre-PHD domain was proposed to stabilize the PHD domains, hence referred to as enhanced PHD (ePHD) domains (**Figure 2E***, full length PHF6)*^29^. To identify the domain(s)/region(s) of PHF6 important for the interaction with RRM2, we generated a series of deletion constructs of the PHF6 coding sequence that were also N-terminally fused to the Nluc donor protein (**Figure 2E**). To validate our experimental set-up, we confirmed fusion protein expression from all transfected constructs (**Figure S2B**). The fragments that did not contain the first PHD domain (Fr1, Fr7 and Fr8) resulted in lower BRET ratios compared to fragments comprising the first PHD domain, except for Fragment 6, which contains only the second PHD domain. Moreover, fragments of the first PHD domain alone (Fr11) or together with the NLS following this domain (Fr10), show approximately the same BRET ratios as full length PHF6, suggesting that PHF6 interacts with RRM2 through its first PHD domain (**Figure 2E**). Previous studies indicated that the two zinc fingers of PHF6 are nearly identical (Todd & Picketts, 2012). Indeed, overlap between the deletion Fragment 6 (containing only the second PHD domain) and Fragment 10 (containing only the first PHD) showed high sequence similarity (73%) (**Figure S2C**). We postulate that, given this high protein sequence similarity, RRM2 can also bind to the second PHD domain, only if it is not structurally hindered by the pre-PHD2 domain, as is the case in Fragment 7 and Fragment 8. Since previous studies revealed that the second PHD domain of PHF6 can bind dsDNA *in vitro* (Liu *et al*, 2014), we propose that the first PHD domain of PHF6 attracts RRM2 towards DNA for replication/repair through its second PHD domain, potentially as part of the previously reported “replitase” complex (Murthy & Reddy, 2006; Técher et al., 2017).

**Figure 2.**
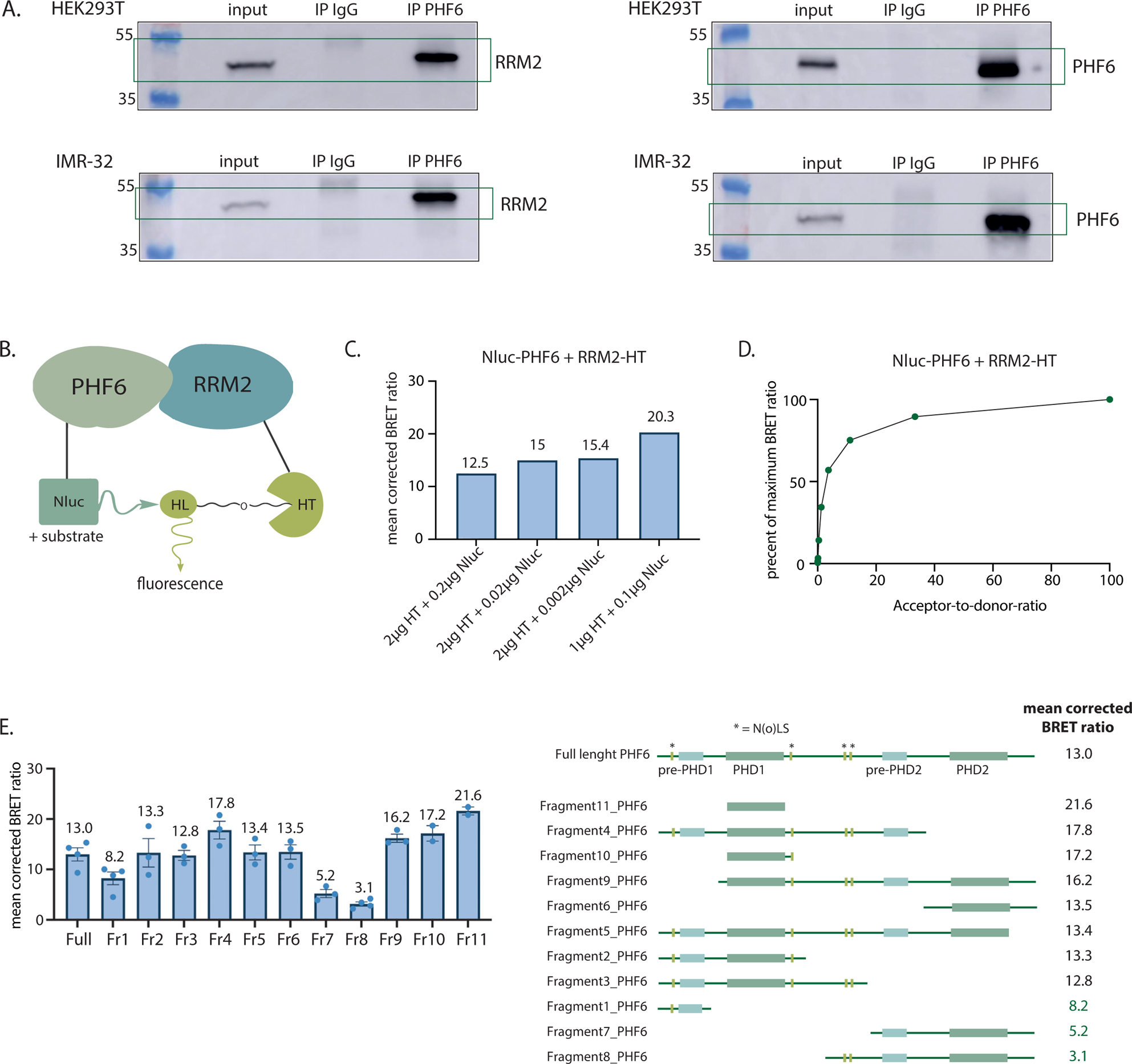
PHF6 interacts with RRM2 through its first PHD domain. **(A)** Immunoblotting for RRM2 (*right*) and PHF6 (*left*) following co-immunoprecipitation analysis using PHF6 as bait in HEK293T (*top*) and IMR-32 (*bottom*) cells. **(B)** Schematic overview of the NanoBRET™ assay for detecting protein-protein interactions by measuring the energy transfer between a NanoLuc® luciferase (Nluc) donor fusion protein with PHF6 and a HaloTag® (HT) acceptor fusion protein with RRM2. Energy transfer only occurs when the fusion proteins are in close proximity to one another (<10nm); **(C)** Mean corrected NanoBRET™ ratios resulting from different amounts of transfected Nluc-PHF6 and RRM2-HT encoding constructs in HEK293T cells; **(D)** Donor saturation assay for the Nluc-PHF6 + RRM2-HT combination results in hyperbolic increase in NanoBRET™ signal; **(E)** (*left*) Mean corrected NanoBRET™ ratios from transfecting 11 different deletion constructs of PHF6 fused to Nluc (Fr1-Fr11) together with the RRM2-HT fusion. Error bars represent the mean ± SEM of 2 to 4 biological replicates (represented by dots) and (*right*) schematic representation of the deletion constructs, sorted from highest to lowest mean NanoBRET™ ratio.

### PHF6 and RRM2 knockdown result in overlapping transcriptional signatures

Our data converge towards RRM2 as a robust novel PHF6 interactor and we previously established a critical role for RRM2 as replicative stress resistance factor in high-risk neuroblastoma (Nunes *et al*, 2022), while the functional role of PHF6 in neuroblastoma tumor development remains elusive. Interestingly, PHF6 is highly expressed in neuroblastoma cell lines (**Figure 3A**) and upregulated during TH-MYCN driven neuroblastoma tumor formation compared to the expression in ganglia from wildtype mice (Weiss *et al*, 1997) (p = 2.43 x 10^−6^) (**Figure 3B**). Moreover, high *PHF6* expression in neuroblastoma patients correlates with poor prognosis (event free and overall survival, p = 2.59 x 10^−8^ and p = 3.62 x 10^−8^ respectively) (**Figure 3C**). In addition, PHF6 expression is increased in high-risk (stage 4) classified neuroblastoma patients and *MYCN*-amplified cases (**Figure 3D**).

**Figure 3.**
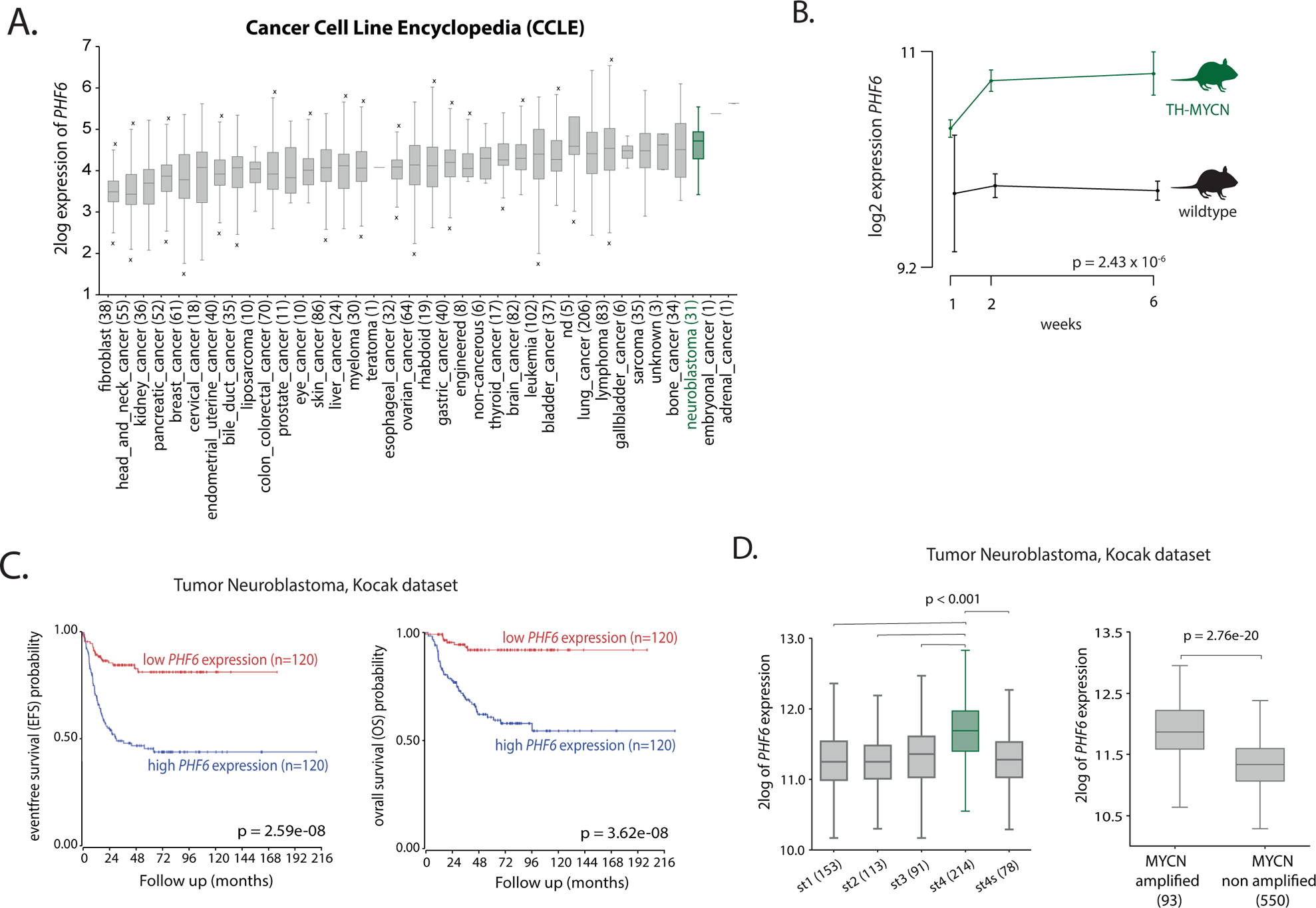
High *PHF6* expression is correlated with poor prognosis in primary neuroblastoma. **(A)** Log_2_ expression of *PHF6* mRNA throughout different cancer cell lines from the ‘Cancer Cell Line Encyclopedia’ (CCLE) database. The boxplots represent the 1st to 3rd quartile of the data values, with the line representing the median value. Whiskers represent values of the outer two quartiles maximized at 1.5 times the size of the box (‘x’ indicates one or more values outside of the whiskers); **(B)** *Phf6* mRNA expression is up-regulated during TH-MYCN–driven neuroblastoma tumor development (De Wyn *et al*, 2021) (p = 2.43 x 10^−6^); **(C)** High *PHF6* expression levels correlate to poor event-free (p = 2.59 x 10^−8^) and overall (p = 3.62 x 10^−8^) neuroblastoma patient survival [Kocak cohort (n = 649); hgserver2.amc.nl, cutoff mode: first_vs_last_quartile]; **(D)** *PHF6* expression is higher in Stage 4 neuroblastoma patients compared to other risk groups (left, p < 0.001) and in *MYCN*-amplified neuroblastomas compared to *MYCN* non amplified cases (right, p = 2.7 x 10^−20^) [Kocak cohort (n = 283); hgserver2.amc.nl].

In view of these findings, we sought to explore whether overlapping molecular responses were observed upon perturbed RRM2 or PHF6 expression in this cellular context. To this end, we selected a panel of four neuroblastoma cell lines to perform transcriptome profiling following PHF6 knockdown. Knockdown of PHF6 was validated through RT-qPCR (**Figure 4A**) and immunoblotting (**Figure 4B**). A robust and overlapping PHF6 knockdown transcriptome signature was identified using ‘Gene set enrichment analysis’ (GSEA) between all four neuroblastoma cell lines (**Figure S3**). Comparison of gene signatures following PHF6 knockdown with our previously published RRM2 knockdown data (GSE161902 (Nunes *et al*, 2022)) revealed significant overlap in direct transcriptional effects as well as with pharmacological RRM2 inhibition using triapine (3AP) (**Figure 4C**). Indeed, common enriched gene sets following RRM2 and PHF6 knockdown were mainly related to amongst others the p53 pathway, G2/M checkpoint, E2F targets and MYC targets (**Figure 4D**).

**Figure 4.**
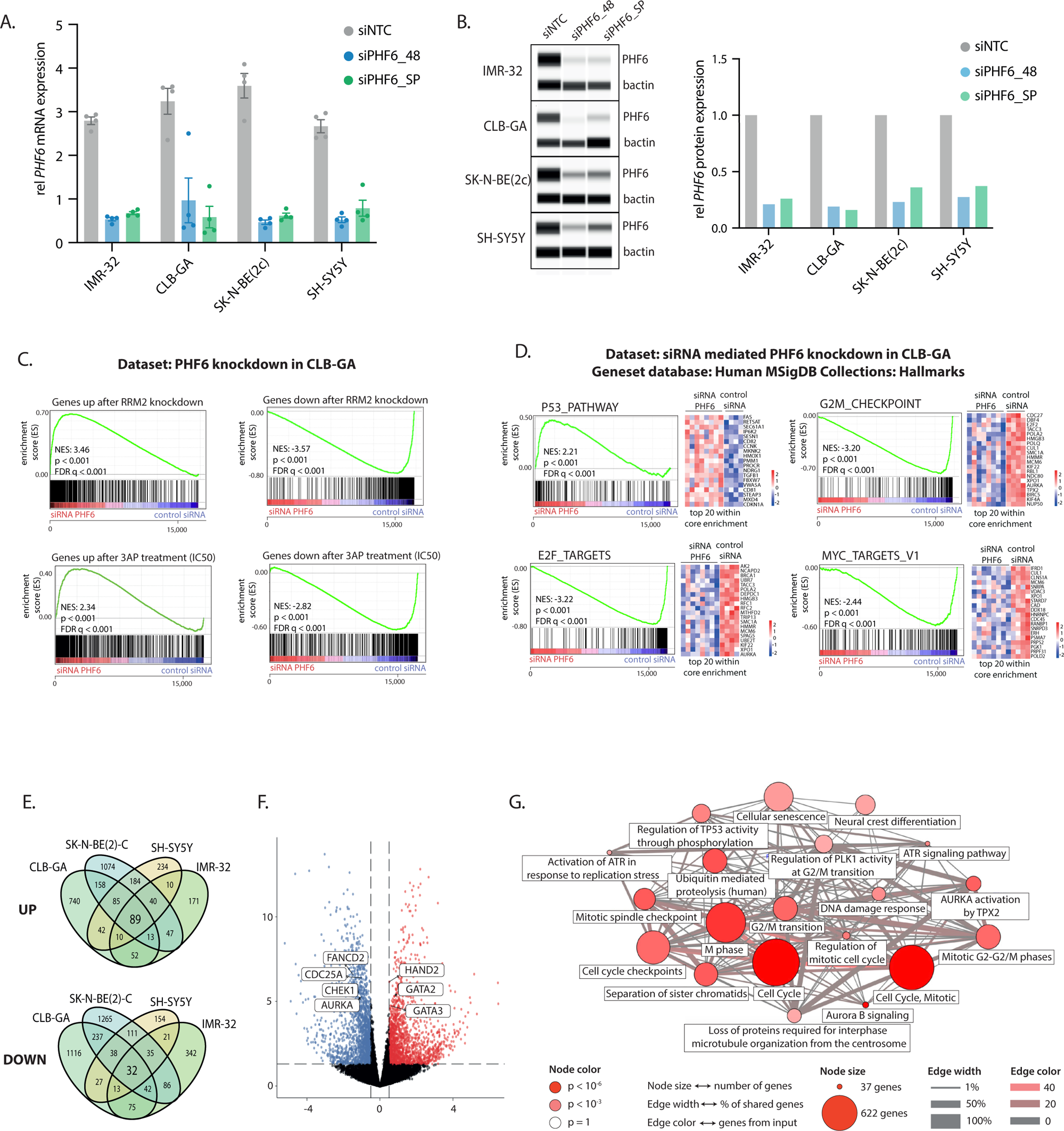
Overlapping gene signatures following knockdown support a functional link between PHF6 and RRM2. **(A)** *PHF6* mRNA expression levels and **(B)** PHF6 protein expression levels upon siRNA mediated PHF6 knockdown in IMR-32, CLB-GA, SK-N-BE(2)-C and SH-SY5Y neuroblastoma cell lines using two different siRNAs (siPHF6_48 and siPHF6_SP) and one no template control siRNA (siNTC). Error bars represent the mean ± SEM of 4 biological replicates (represented by dots); **(C)** Gene set enrichment analysis (GSEA) shows a significant overlap between up- and down regulated genes upon PHF6 knockdown using siRNAs and genes downregulated following either RRM2 knockdown pharmacological RRM2 inhibition using 3AP (GSE161902 (Nunes *et al*, 2022)); **(D)** GSEA of the RNA-seq based transcriptome profiles after PHF6 knockdown in CLB-GA cells using the Hallmark MSigDB gene sets **(E)** Venn diagram showing the overlap of up- and downregulated genes identified by RNA-sequencing upon PHF6 knockdown in CLB-GA, SK-N-BE(2)-C, SH-SY5Y and IMR-32 neuroblastoma cells compared to the siRNA control; **(F)** Volcano plot displaying significantly upregulated (red) and downregulated (blue) genes upon siRNA mediated knockdown of PHF6 in CLB-GA cells; **(G)** ConsensusPathDB over-representation analysis of downregulated genes upon PHF6 knockdown showing enrichment in amongst others DNA damage response and cell cycle checkpoint regulation processes.

Next, overlap between the significantly differentially expressed genes (DEG) (adjusted P-value < 0.05) upon PHF6 knockdown in at least two out of the four cell lines revealed upregulation of 730 genes and downregulation of 717 genes (**Figure 4E**). Interestingly, genes involved in replication and replication stress response pathways comprising *FANCD2*, *CDC25A*, *CHK1* and *AURKA* were downregulated (**Figure 4F**). Enrichment analysis using the ConsensusPathDB (Kamburov *et al*, 2009) tool for the 717 common down regulated genes, shows enrichment for genes involved in G2/M transition, ATR signaling, DNA damage responses and cell cycle regulation (**Figure 4G**).

### PHF6 knockdown induces replicative stress and DNA damage in neuroblastoma cells

Given the recently described role for PHF6 as important factor in fork restart after DNA damage (Warmerdam *et al*, 2019) and the above data suggesting a link between PHF6 and replicative stress, we hypothesized that elevated PHF6 levels could contribute to alleviating replicative stress levels. In line with previous studies in B-ALL (Feliciano *et al*, 2017), knockdown of PHF6 did not significantly affect neuroblastoma cell confluence as measured by Incucyte live cell imaging (**Figure 5A**). Interestingly, accordingly to what we have previously shown for RRM2 (Nunes *et al*, 2022), pCHK1^S345^, pRPA32 and γH2AX levels were elevated upon knockdown of PHF6, indicative of replicative stress and DNA damage induction (**Figure 5B**). Previous studies reported CHK1 as a synthetic lethal target in *MYCN*-amplified neuroblastoma and neuroblastoma was ranked as the most sensitive pediatric cancer for CHK1 inhibition using prexasertib (Cole *et al*, 2011). Therefore, we scored publicly available gene signatures related to prexasertib sensitivity and resistance in a large cohort of primary neuroblastoma patients (GSE62564) (Blosser *et al*, 2020), revealing that low *PHF6* expression correlates with prexasertib resistance (p = 1.25 x 10^−7^) while high *PHF6* expression levels are correlated with sensitivity for prexasertib treatment (p < 2 x 10^−16^), indicating ATR-CHK1 dependency in neuroblastoma tumor cells with high PHF6 expression (**Figure 5C**).

**Figure 5.**
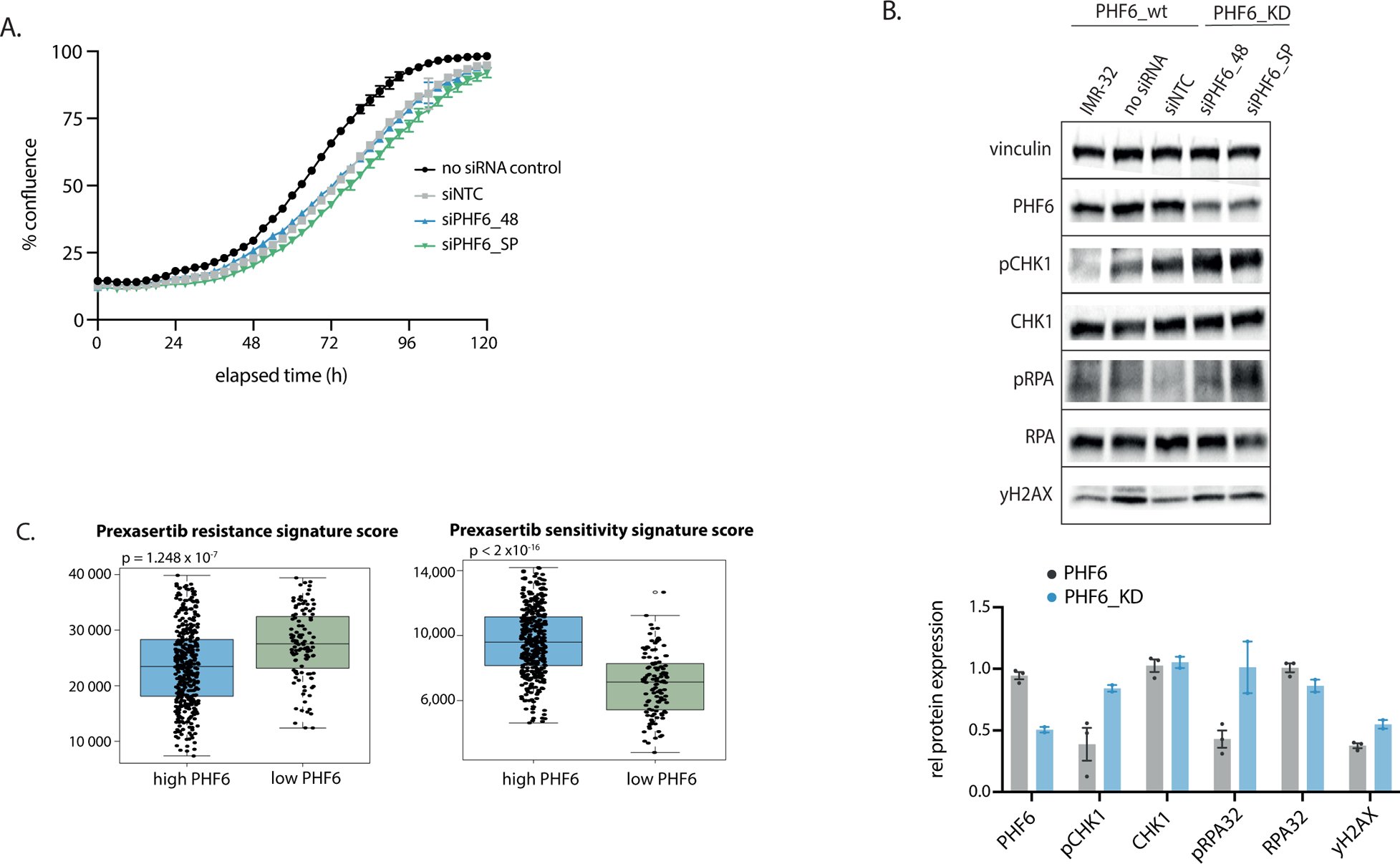
High PHF6 expression leads to increased replicative stress. **(A)** Time-resolved monitoring of cell confluence following siRNA mediated PHF6 knockdown in neuroblastoma cells using Incucyte live cell imaging; **(B)** Immunoblotting for various DNA damage markers in IMR-32 cells upon siRNA mediated knockdown of PHF6 (*top*), quantification of the blot using ImageJ is shown (*bottom*) panel; **(C)** Signature score analysis of prexasertib sensitivity and resistance gene signatures in a large primary cohort of neuroblastoma cases (GSE62564).

### PHF6 binds to acetylated histone 3 at lysine 53 (H3K56ac) at regions of DNA damage and fork stalling

Enrichment analysis using the ENRICHR (Xie *et al*, 2021) tool for the 717 common down regulated genes upon PHF6 knockdown in at least two out of the four neuroblastoma cell lines also revealed overlap with amongst others H3K9ac and H3K56ac ChIP-seq marked target genes (**Figure 6A**), both previously identified as DNA-damage responsive histone modifications (Tjeertes et al., 2009). Interestingly, H3K9ac has been implicated in regulating transcriptional repression following DNA damage, mediated by CHK1 (Shimada *et al*, 2008). In contrast, H3K56ac is a histone modification marking nascent chromatin or sites of replication fork stalling (Whale *et al*, 2022), but is also deposited at sites of transcriptional activation (Williams et al., 2008; Yuan et al., 2009). To test this, co-IP experiments were performed and indeed showed PHF6 binding with H3K56ac, while no evidence for binding to H3K9ac was found. We also observed slightly increased binding upon low-dose etoposide treatment (0.1µM) in IMR-32 cells (**Figure 6B**), in keeping with our proposed role for PHF6 in DNA replication and repair. Acetylation of H3K56 results from amongst others CBP/EP300 activity predominantly at highly transcribed regions (Das *et al*, 2009). In view of our finding that PHF6 binds to H3K56ac, we performed CUT&RUN to map PHF6 and H3K56ac sites in IMR-32 and CLB-GA neuroblastoma cells and observed overlap between the binding sites for PHF6 and H3K56ac in both cell lines (**Figure 6C**). To further investigate the role of PHF6 in relation to dsDNA breaks and stalled replication forks, we performed additional CUT&RUN analyses for fork associated proteins BRCA1, DDX11, ATR and FANCD2 and double strand DNA damage marker γH2AX in IMR-32 cells. Also, for these markers significant overlap with PHF6 bound regions was noted (**Figure 6D**). These results support the critical role of PHF6 at the crossroad of DNA damage and replicative stress responses.

**Figure 6.**
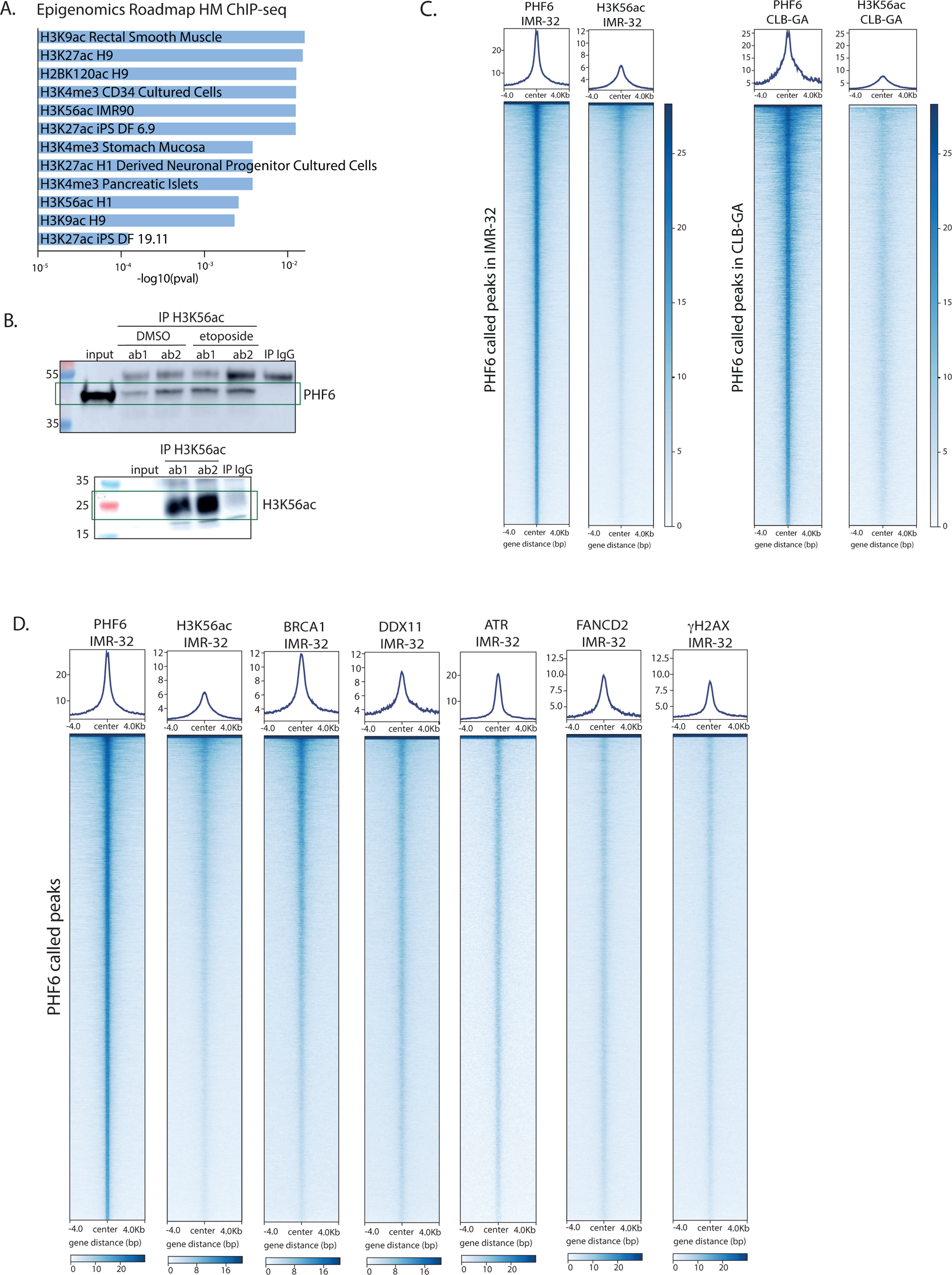
PHF6 binds to the H3K56ac chromatin mark at regions of DNA damage and fork stalling. **(A)** Enrichment analysis using ENRICHR of the downregulated genes upon PHF6 knockdown shows enrichment for ChIP-sequencing identified targets of H3K27ac, H3K4me3, H3K9ac and H3K56ac associated genes; **(B)** Immunoblotting for PHF6 (*top*) and H3K56ac (*bottom*) after co-immunoprecipitation analysis with H3K56ac as using two different antibodies (ab1 and ab2) in IMR-32 neuroblastoma cells upon DMSO and etoposide treatment (0.1µM). **(C)** Heatmap profiles −4kb and +4kb around the summit of PHF6 CUT&RUN peaks showing the significant overlap (min overlap = 10 bp) between PHF6 and H3K56ac CUT&RUN peaks in IMR-32 (Fisher’s Exact Test p-value < 2.2 x 10^−16^) and CLB-GA (Fisher’s Exact Test p-value < 2.2 x 10^−16^) cells. Peaks were ranked according to the sums of the peak scores across the datasets in the heatmap; **(D)** Heatmap profiles −4kb and +4kb around the summit of PHF6 CUT&RUN peaks, representing overlap of PHF6 CUT&RUN peaks in IMR-32 cells with BRCA1, DDX11, ATR, FANCD2 and gH2AX. Peaks were ranked according to the sums of the peak scores across all datasets in the heatmap.

### The PHF6 chromatin binding landscape overlaps with adrenergic core regulatory circuitry transcription factor binding in neuroblastoma

Next, we performed enrichment analysis using ENRICHR (Xie *et al*, 2021) on the overlapping binding sites between PHF6 and H3K56ac. This revealed significant enrichment of ChIP-seq identified targets of CTCF and FOXM1 as well as ISL1, PHOX2B, GATA3, HAND2, MYCN and TBX2, which are members of the previously described adrenergic neuroblastoma ‘core regulatory circuitry’ (CRC) (Boeva *et al*, 2017; Van Groningen *et al*, 2017; Shendy *et al*, 2022). Notably, also EP300 ChIP-seq targets were found to be enriched in the set of overlapping sites (**Figure 7A**). Neuroblastoma tumors display cellular heterogeneity, comprising both adrenergic and neural crest like (mesenchymal) cell types. Both cell identities are characterized by super-enhancer marked transcription factors, which are controlling the gene expression program of the two neuroblastoma cell identities (Shendy *et al*, 2022). Interestingly, among the 730 common upregulated genes after siRNA mediated PHF6 knockdown in two out of the four neuroblastoma cell lines, we found *GATA2*, *GATA3* and *HAND2* (**Figure 4F**). Using the ENRICHR tool (Xie *et al*, 2021) for enrichment analysis on the 717 common down regulated genes, we also found a significant overlap with ChIP-seq identified targets of E2F1, FOXM1 and different adrenergic CRC transcription factors, including ISL1, PHOX2B, GATA3, HAND2, MYCN and TBX2 (**Figure 7B**). Moreover, motif analysis using HOMER also revealed significant enrichment for CTCF and adrenergic CRC transcription factor motifs of ASCL1, PHOX2A, ISL1, PHOX2B, HAND2, GATA3 and GATA2 in the PHF6 binding sites (**Figure 7C**), further supporting significant overlap between PHF6 and the CRC transcription factors in neuroblastoma. Indeed, when comparing ChIP-seq data from the studies of Boeva *et al*. (Boeva *et al*, 2017) and Durbin *et al*. (Durbin *et al*, 2018), we observed significant overlap in the binding sites of PHF6 with HAND2, PHOX2B, GATA3, ASCL1 and MYCN, predominantly at the PHF6 enhancer associated peaks (**Figure 7D**). As these CRC members are known to be highly transcribed in neuroblastoma, these regions are inherently at higher risk to suffer from replicative stress due to head-on collisions between the replication and transcription machinery (Shendy *et al*, 2022). Therefore, we propose that PHF6 is recruited to these regions, possibly through the presence of H3K56ac positive collapsed replication bubbles (Whale *et al*, 2022), attracting RRM2 to relief transcription-associated DNA damage. Indeed, binding of PHF6 and H3K56ac was confirmed at the *PHOX2B*, *GATA3*, *HAND2* and *MYCN* loci (**Figure 7E**).

**Figure 7.**
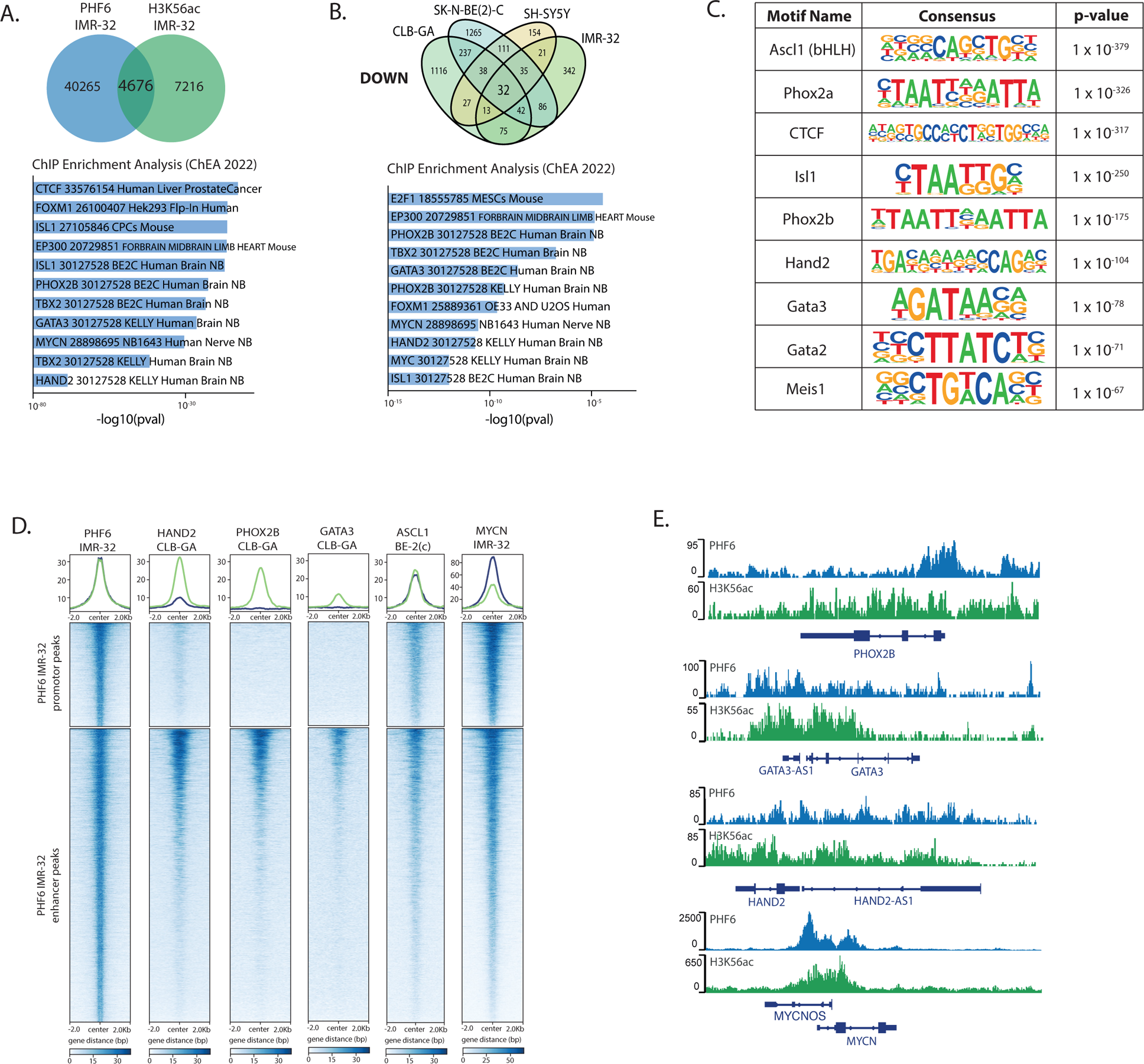
CUT&RUN reveals binding of PHF6 to members of the adrenergic CRC in neuroblastoma. **(A)** Venn diagram (*top*) showing overlap between PHF6 and H3K56ac CUT&RUN peaks in IMR-32 cells and enrichment analysis (*bottom)* using ENRICHR of the PHF6 and H3K56ac overlapping peaks, showing enrichment for ChIP-seq identified targets of CTCF, FOXM1 and ADR CRC members ISL1, PHOX2B, HAND2, TBX2, GATA3 and MYCN; **(B)** Venn diagram showing the overlap of downregulated genes identified by RNA-sequencing upon PHF6 knockdown in CLB-GA, SK-N-BE(2)-C, SH-SY5Y and IMR-32 neuroblastoma cells compared to the siRNA control (*top*) and enrichment analysis using ENRICHR of the downregulated genes upon PHF6 knockdown shows enrichment (*bottom)* for ChIP-sequencing identified targets of E2F1, FOXM1 and neuroblastoma adrenergic core regulatory circuitry members (ADR CRC) including ISL1, PHOX2B, GATA3, HAND2, MYCN and TBX2; **(C)** Homer motif analysis of the PHF6 CUT&RUN peaks in IMR-32 cells; **(D)** Heatmap profiles −2kb and +2kb around the summit of PHF6 CUT&RUN peaks, representing overlap of PHF6 CUT&RUN peaks in IMR-32 cells with MYCN CUT&RUN peaks as well as publicly available ChIP-seq peaks of HAND2, PHOX2B and GATA3 in CLB-GA [GSE90683], ASCL1 in SK-N-BE(2)-C [GSE159613] which were grouped for promoters or enhancers (using homer annotation), and ranked according to the sums of the peak scores across all datasets in the heatmap; **(E)** PHF6 and H3K56ac binding activity at the PHOX2B, GATA3, HAND2 and MYCN loci in IMR-32 cells. Signals represent log likelihood ratio for the CUT&RUN signal compared to control signal (RPKM normalized). All peaks are called by MACS2 (q < 0.05).

Previously, we performed CasID experiments in SK-N-BE(2)-C to identify upstream regulators of *RRM2* expression (Nunes *et al*, 2022). This revealed that PHF6 as well as NuRD and PAF1 protein complex members, both previously described as PHF6 interactors (Todd & Picketts, 2012; Zhang et al., 2013), are binding to the *RRM2* promotor (**Figure S4A, *top***). In light of this finding, we also performed CUT&RUN for PHF6 in SK-N-BE(2)-C, and confirmed binding of PHF6 to the *RRM2* promotor in all three cell lines that we tested (**Figure S4A, *bottom***). In addition, we observed strong overlap between the PHF6 binding sites (**Figure S4B**), resulting in 3171 genes that are bound by PHF6 in all three cell lines (**Figure S4C, *top***). ENRICHR (Xie *et al*, 2021) analysis on these overlapping PHF6 binding sites revealed significant enrichment for binding sites of FOXM1, CTCF and those of the adrenergic CRC transcription factors PHOX1B, ISL1, MYCN, GATA3, TBX2, amongst others (**Figure S4C, *bottom*)**. Peak annotation distribution analysis shows that PHF6 mainly binds to intron and intergenic regions (**Figure S4D, *top***), which are significantly overlapping with active promotor/enhancer histone marks (**Figure S4D, *bottom* and S4E**). Moreover, we also observed strong overlap between the binding profiles of PHF6 and differentially expressed genes after siRNA mediated PHF6 knockdown, showing gene expression regulation at PHF6 bound regions (**Figure S4F)**

## Discussion

Rapidly dividing cancer cells are more prone to elevated replication stress levels, resulting from increased replication fork stalling. This may be triggered amongst others through nucleotide depletion, replication-transcription conflicts and R-loop formation. Replication stress occurs in normal cells mainly during early S-phase (Buisson *et al*, 2015) and can drive tumor initiation and progression, but can also impede tumor growth due to accumulation of excessive DNA damage. In response to these stress levels, cancer cells upregulate DNA damage signaling and the expression of factors enabling to tolerate replication stress. Previous studies (Gaggioli *et al*, 2023) support an important role for the epigenetic machinery in DNA repair processes and chromatin dynamic re-organization when DNA replication forks are challenged. In this context, the hitherto presumed chromatin reader PHF6 has been identified as a major replication fork restart factor (Warmerdam *et al*, 2019). This is in line with earlier observations that PHF6 is a substrate for ATM and ATR phosphorylation (Matsuoka *et al*, 2007) and more recently as R-loop modulator (Yan *et al*, 2022), the latter being recognized as a potent source of replication stress (Saxena & Zou, 2022). It remains enigmatic thus far how PHF6 functionally contributes to replication fork restart and whether its association with chromatin is involved in this process. Using an unbiased approach, we identified RRM2 as robust PHF6 interactor in both normal and malignant cell types, suggesting that this interaction has a generic function in cells. Through NanoBRET^TM^ based deletion mapping, we could identify the critical interface for the physical interaction of PHF6 with RRM2 to the first PHD domain of PHF6. Given that RRM2 is the catalytic and rate-limiting factor of the RRM1-RRM2 heterodimer, referred to as the ribonucleotide reductase enzyme, we postulate that the interaction of RRM2 with PHF6 installs a functional replitase (Técher *et al*, 2017) which has been proposed for a long time, but remains thusfar unrecognized.

To further understand the role of the PHF6-RRM2 interaction, we selected neuroblastoma cells which exhibit strong expression levels of both proteins and renown for high intrinsic *MYCN* oncogene induced replication stress. Differential gene expression analyses following PHF6 knockdown hinted towards a possible functional connection with histone H3 lysine 56 acetylation. Through further co-IP analysis, we could indeed reveal PHF6 as a H3K56ac reader. This chromatin modification is found at sites of nascent DNA, linked to suppression of DNA polymerase activity and elongation during break-induced DNA repair (Che *et al*, 2015) and controls gene expression homeostasis during S-phase (Topal *et al*, 2019). Recent work from the Houseley team pointed out that H3K56ac sites are enriched in the coding sequence of highly transcribed genes due to collapse of replication bubbles forming so-called H3K56ac ‘scars’ (Whale *et al*, 2022). Previously, H3K56ac was also shown to be a critical factor required for cell cycle re-entry following DNA repair (Battu et al., 2011), preferentially deposited in nucleosomes with a short turn-over (Masumoto *et al*, 2005). In this respect, we speculate that PHF6 facilitates and controls high transcriptional rates of super-enhancer associated genes in cancer cells and that PHF6 chromatin binding, through association with H3K56ac, is required to ensure finalization of DNA repair and subsequent release from checkpoint inhibition in response to replication stress. In line with these findings, PHF6 binding sites also overlap with γH2AX, FANCD2, DDX11 and BRCA1 bound chromatin regions. This observation suggests a prominent function for PHF6 in DNA damage repair and fork protection under stress (Mahtab *et al*, 2021; Qiu *et al*, 2021). Previous studies showed that BRCA1 is recruited by MYCN to promoter-proximal regions to prevent MYCN-dependent accumulation of stalled RNA polymerase II (RNAPII) (Herold *et al*, 2019). Moreover, BRCA1 was also reported to regulate *RRM2* expression (Rasmussen *et al*, 2016). Furthermore, the Ferrando team suggested recently that PHF6 is involved in nuclear retention of BRCA1 upon DNA damage, with direct implications in homologous recombination mediated DNA repair (Warmerdam *et al*, 2019). Interestingly, DDX11 also acts downstream of 53BP1 to mediate HR repair (Jegadesan & Branzei, 2021). Furthermore, our CUT&RUN analysis also revealed for the first time that genome-wide binding sites of PHF6 significantly overlap with those for previously described neuroblastoma specific CRC factors including MYCN, HAND2, PHOX2B, GATA3 and ISL1. In addition, these binding sites were strongly enriched at H3K4me1/H3K27ac marked active enhancers.

Our IP-MS experiments also displayed enrichment for other putative relevant factors in these processes. Amongst others we found a consistent enrichment for the chromatin interacting protein and DNA damage responsive factor PATZ1. Although its functional role in pediatric cancer remains enigmatic thus far, PATZ1 has been shown to be overexpressed as fusion transcript with EWSR1 in glioma (Siegfried *et al*, 2019), while a genome-scale CRISPR-Cas9 screen in *MYCN* amplified neuroblastoma cells revealed dependency of neuroblastoma on PATZ1 expression as well (Durbin *et al*, 2018). Other recurrent PHF6 interacting hits in our IP-MS experiments included TOP2B that is recruited to stalled forks (Tian *et al*, 2021), ABCF1 described as sensor of genomic integrity (Choi *et al*, 2021) and two components of the nuclear pore complex (NPC), *i.e*. RANBP2, a SUMO E3 ligase and ELYS, an important factor for assembly of the NPC. Interestingly, these latter two factors also functionally connect to replicative stress resistance (Franz et al., 2007; Sarangi & Zhao, 2015; Xiao et al., 2015). The NPC is a very dynamic structure. In addition to its role as gateway between the nucleoplasm and cytoplasm, it actively participates in regulating replication fork dynamics, protection to replicative stress and DNA repair (Kuhn & Capelson, 2019; Whalen et al., 2020).

In summary, we propose PHF6 as a key novel chromatin adaptor protein in functional concertation with RRM2 under replicative stress and as component of the earlier proposed ‘replitase’ (Técher *et al*, 2017), implicated in active regulation of RRM2 function during replication and DNA repair (**Figure 8**). In this context, PHF6 potentially brings RRM2 as essential rate-limiting factor for dNTP production in the immediate vicinity of these sites, requiring unperturbed DNA synthesis in order to allow for uninterrupted S-phase transition and protect rapidly dividing cancer cells from toxic accumulation of transcription-coupled DNA damage.

**Figure 8.**
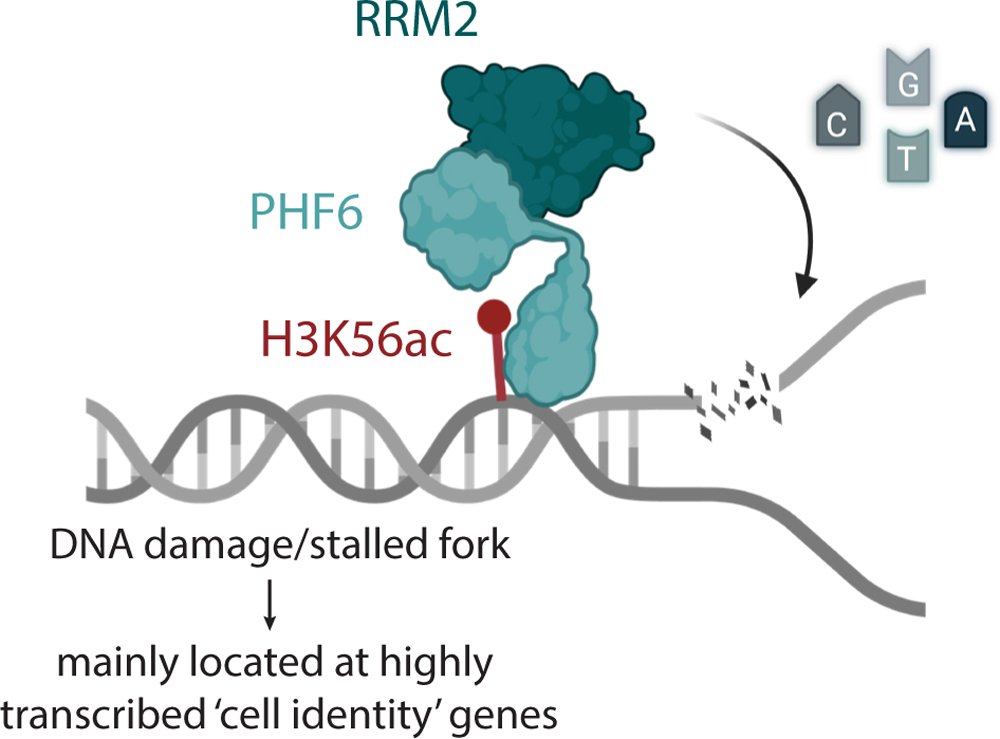
PHF6 directs RRM2 to regions of stalled replication forks and DNA damage to relieve transcription-associated DNA damage. We hypothesize that PHF6 is a crucial component of the so-called ‘replitase’ complex to direct RRM2 to regions of stalled replication forks or DNA damage, mediated by binding to the DNA damage related H3K56ac histone mark with a potential role in suppressing transcription-associated DNA damage, mainly at highly transcribed, so-called cell identity genes.

## Materials and methods

### Cell culture

The human neuroblastoma cell lines (CLB-GA, IMR-32, SH-SY5Y and SK-N-BE(2)-C), the retinal pigmented epithelial cell line (RPE) and human fetal kidney–derived cell line (HEK293T) were grown in RPMI 1640 supplemented with 10% fetal calf serum (FCS), penicillin/ streptomycin (100 IU/ml), and 2 mM L-glutamine. The leukemia cell line ALL-SIL was grown in RPMI 1640 supplemented with 20% fetal calf serum (FCS), penicillin/ streptomycin (100 IU/ml), and 2 mM L-glutamine. HCT116 cells were cultured in McCoy’s 5A medium also supplemented with 10% fetal calf serum (FCS), penicillin/ streptomycin (100 IU/ml), and 2 mM L-glutamine. All cell lines used were cultured in 5% CO2 atmosphere at 37°C on plastic cultured plates. Short Tandem Repeat (STR) genotyping was used to validate cell line authenticity prior to performing the described experiments and Mycoplasma testing was done on a monthly basis. Origin of the cell lines and products used for cell culture can be found in **Table S1** and **Table S2** respectively.

### Co-immunoprecipitation followed by mass-spectrometry or Western Blot analysis

Co-immunoprecipitation (co-IP) with PHF6 as bait protein was performed by lysing 10M cells/sample in 1ml cold radioimmunoprecipitation assay (RIPA) buffer (12.7 mM; 150 mM NaCl, 50 mM Tris-HCl (pH 7.5), 0.01% SDS solution, and 0.1% NP-40) supplemented with protease and phosphatase inhibitors. Lysates were incubated with 2µg antibody (PHF6 or human IgG) for 4 hours. After adding 20 µl Protein A UltraLink® Resin (Life Technologies, 53139), the mixture was rotated overnight at 4°C.

After the overnight incubation, peptides were re-dissolved in 20 µl loading solvent A (0.1% TFA in water/ACN (98:2, v/v)) of which 2 µl was injected for LC-MS/MS analysis on an Ultimate 3000 RSLC nano LC (Thermo Fisher Scientific, Bremen, Germany) in-line connected to a Q Exactive mass spectrometer (Thermo Fisher Scientific). Trapping was performed at 10 μl/min for 4 min in loading solvent A on a 20 mm trapping column (made in-house, 100 μm internal diameter (I.D.), 5 μm beads, C18 Reprosil-HD, Dr. Maisch, Germany). The peptides were separated on a 25 cm in needle packed column (made in-house, 75 μm internal diameter (I.D.), 3 μm beads, C18 Reprosil-HD, Dr. Maisch, Germany) kept at a constant temperature of 50°C). For ALL-SIL, CLB-GA and IMR-32, peptides were eluted by a non-linear gradient starting at 2% MS solvent B reaching 55% MS solvent B (0.1% FA in water/acetonitrile (2:8, v/v)) in 123 min, 99% MS solvent B in 12 minutes followed by a 6-minute wash at 70% MS solvent B and re-equilibration with MS solvent A (0.1% FA in water). For the other cell lines, peptides were eluted by a linear gradient reaching 30% MS solvent B (0.1% FA in water/acetonitrile (2:8, v/v)) after 74 (SK-N-BE(2)-C, SH-SY5Y and HCT116)/ 104 (HEK293T and RPE, 50% MS solvent B after 99 min (SK-N-BE(2)-C, SH-SY5Y and HCT116) / 121 min (HEK293T and RPE) and 99% MS solvent B at 100 min (SK-N-BE(2)-C, SH-SY5Y and HCT116) / 125 min (HEK293T and RPE), followed by a 5-minutes wash at 99% (SK-N-BE(2)-C, SH-SY5Y and HCT116) / 70% (HEK293T and RPE) MS solvent B and re-equilibration with MS solvent A (0.1% FA in water). The first 9 min the flow rate was set to 750 nl/min after which it was kept constant at 300 nl/min. The mass spectrometer was operated in data-dependent mode, automatically switching between MS and MS/MS acquisition for the 5 (IMR-32) / 10 (other cell lines) most abundant ion peaks per MS spectrum. Full-scan MS spectra (400-2000 m/z) were acquired at a resolution of 70,000 in the Orbitrap analyzer after accumulation to a target value of 3,000,000. The 5 (IMR-32) / 10 (other cell lines) most intense ions above a threshold value of 13,000 (IMR-32) / 17,000 (other cell lines) were isolated with a width of 2 m/z for fragmentation at a normalized collision energy of 25% after filling the trap at a target value of 50,000 for maximum 80 (IMR-32) / 60 (other cell lines). MS/MS spectra (200-2000 m/z) were acquired at a resolution of 17,500 in the Orbitrap analyzer.

Analysis of the mass spectrometry data was performed in MaxQuant (version 2.2.0.0) with mainly default search settings including a false discovery rate set at 1% on PSM, peptide and protein level. Spectra were searched against the human reference proteome (version of 01_2023, UP000005640). The mass tolerance for precursor and fragment ions was set to 4.5 and 20 ppm, respectively, during the main search. Enzyme specificity was set as C-terminal to arginine and lysine, also allowing cleavage at proline bonds with a maximum of two missed cleavages. Variable modifications were set to oxidation of methionine residues and acetylation of protein N-termini. Matching between runs was enabled with a matching time window of 0.7 minutes and an alignment time window of 20 minutes. Only proteins with at least one unique or razor peptide were retained. Proteins were quantified by the MaxLFQ algorithm integrated in the MaxQuant software. A minimum ratio count of two unique or razor peptides was required for quantification. Further data analysis of the results was performed with an in-house R script, using the proteinGroups output table from MaxQuant. Reverse database hits were removed, LFQ intensities were log2 transformed and replicate samples were grouped. Proteins with less than three valid values in at least one group were removed and missing values were imputed from a normal distribution centered around the detection limit (package DEP (Zhang *et al*, 2018)). To compare protein abundance between pairs of sample groups (IgG vs PHF6 sample groups), statistical testing for differences between two group means was performed, using the package limma (Ritchie *et al*, 2015). Statistical significance for differential regulation was set to a false discovery rate (FDR) of <0.05 and|log2FC| ≥ 0.6. The mass spectrometry proteomics data have been deposited to the ProteomeXchange Consortium via the PRIDE (Perez-Riverol *et al*, 2022) partner repository with the dataset identifier PXD041025 and 10.6019/PXD041025.

For the co-IP based validation of the PHF6-RRM2 interaction through immunoblotting, the mixtures were centrifugated, after overnight incubation on 4°C, for 2min at 2000 rpm and washed in RIPA buffer three times. The samples were denaturated in 1X Laemli denaturation buffer supplemented with β-mercaptoethanol and incubated at 95°C for 10minutes. The denatured samples were loaded on 10% SDS–polyacrylamide gel electrophoresis (SDS-PAGE) gels with 10× Tris/Glycine/SDS buffer and run for 1 hour at 100 V, blotted on nitrocellulose membranes in 10% of 10× Tris/Glycine buffer for 1 hour at 100 V. The membranes were blocked during 1 hour in 5% milk in Tris-buffered saline with 0.1% Tween® 20 detergent (TBST). Primary antibody incubations with the RRM2, H3K56ac or PHF6 antibody (**Table S3**) were done in blocking buffer overnight at 4°C. Blots were washed three times with TBST before the incubation for 1 hour with Veriblot secondary antibodies. The immunoblots were visualized by the enhanced chemiluminescent West Dura or West Femto (Bio-Rad) using the Amersham Imager 680. Origin of products used for Western Blotting can be found in **Table S2**, antibodies used can be found in **Table S3**.

### NanoBRET™ analysis

The NanoBRET™ Nano-Glo detection System (Promega) was used according to the manufacturer’s instructions. In short, expression plasmids with RRM2 or PHF6 fused to HaloTag® or NanoLuc® at N or C terminal site, were made using the NanoBRET™ PPI MCS Starter System (Promega, cat n° N1811). The RRM2 fragments were amplified by PCR using Phusion polymerase (NEB, M0530L) and OriGene clone GC-Z9335-GS as template. For PHF6 fragments, OriGene clone GC-Z2633-CF served as template. All fragments and vectors pNLF1-C (CMV/Hygro), pNLF1-N (CMV/Hygro), pHTN HaloTag® CMV-neo and pHTC HaloTag® CMV-neo were digested with restriction enzymes and subsequently ligated (Takara, cat n°6023). Primers and corresponding restriction digests can be found in **Table S4**. The sequences of the constructed plasmids were verified by Sanger DNA sequencing (Eurofins Genomics, SupremeRun Tube Service) and the overexpression in HEK293T was validated using a Simple Western™ automated blot system (WES™ system, ProteinSimple, Bio-Techne) according to manufacturer’s instructions. In total, 1.2 µg of protein lysate was used, primary antibodies anti-PHF6, anti-RRM2 and anti-NanoLuc® antibody were used (see **Table S3** for antibody information). Next, HEK293T cells were co-transfected with HaloTag and Nluc fusions using FuGENE reagent (Promega). After 20h, the transfected cells were trypsinized, washed with PBS, resuspended in assay medium (Opti-MEM I Reduced Serum Medium, no phenol red, supplemented with 4% FBS), re-plated to a 96-well plate, and incubated overnight with HaloTag NanoBRET™ 618 Ligand (100 nmol/L final concentration) or DMSO as negative control. NanoBRET™ Nano-Glo Substrate was added, donor emission (460 nm) and acceptor emission (618 nm) signals were measured using the GloMax Discover System (Promega). Raw NanoBRET™ ratio values with milliBRET units (mBU) were calculated as RawBRET = (618 nm Em/460 nm Em) × 1000. Corrected NanoBRET™ ratio values with milliBRET units were calculated as corrected BRET = RawBRET of experimental sample – RawBRET of negative control sample, which reflected energy transfer from the bioluminescent protein donor to the fluorescent protein acceptor reflecting the protein-protein interaction.

### siRNA-mediated knockdown of PHF6

IMR-32, CLB-GA, SH-SY5Y and SK-N-BE(2)-C neuroblastoma cells were transfected using the Neon transfection system (MPK10096) with two siRNAs toward PHF6: siPHF6-48 (s38848, Life technologies, #439242) and siPHF6-SP (SMARTPool 84295, Dharmacon, L-014862-00-0005). As negative control, a scrambled siRNA (Ambion, #AM4635) was used. After transfection, cells were seeded in a T-25 flask at a density of 2 x 10^6^ cells per flask. For proliferation analysis, 14000 cells/well were seeded in a 96-well plate, treated with either DMSO or 3AP and analyzed using the IncuCyte^TM^ Live Cell Imaging System. Each image was analyzed through the IncuCyte Software and cell proliferation was determined by the area (percentage of confluence) of cell images over time. The cells seeded in a T-25 flask were collected for RNA and protein isolation, 48 hours after transfection. Knockdown of PHF6 was evaluated by RT-qPCR and immunoblotting.

### Western Blotting

Cells were lysed in RIPA supplemented with protease and phosphatase inhibitors and rotated for 1 hour at 4°C. The lysates were cleared by centrifugation at 10,000 rpm in a microcentrifuge for 10 min at 4°C. Concentrations were determined using the Pierce BCA Protein Assay Kit. The lysates were denaturized before loading on a gel, by adding 5X Laemli denaturation buffer supplemented with β-mercaptoethanol. 30µg of protein extracts were loaded on SDS-PAGE gels with 10× Tris/Glycine/SDS buffer and run for 1 hour at 100 V. Samples were blotted on nitrocellulose or polyvinylidene difluoride (PVDF) membranes in 1x Tris/Glycine buffer and 20% of methanol. The membranes were blocked during 1 hour in 5% nonfat dry milk or 5% BSA in TBST. Primary antibody incubations for PHF6, pCHK1(Ser345), CHK1, pRPA32, RPA32, γH2AX, β-actin and vinculin were done in blocking buffer overnight at 4°C. Blots were washed three times with TBST and incubated for 1 hour with secondary antibodies. The immunoblots were visualized by using the enhanced chemiluminescent WES Dura or WES Femto (Bio-Rad) using the Amersham Imager 680. Quantification was performed through ImageJ software, where the area from each protein was normalized to the loading protein in respect to each blot. Products used for Western Blotting can be found in **Table S2**, antibodies can be found in **Table S3.**

### RNA-seq and RT-qPCR validation

RNA was isolated using the miRNeasy kit from QIAGEN including an on-column deoxyribonuclease treatment. Concentrations were measured using NanoDrop (Thermo Fisher Scientific). Libraries for mRNA-seq were prepared using the QuantSeq 3ʹ mRNA library prep kit (Lexogen, Vienna, Austria) with UMI barcoding according to the manufacturer’s protocol. The libraries were equimolarly pooled and sequenced on an Illumina NextSeq 500 high-throughput flow cell, generating single-end 75-bp reads. Adequate sequencing quality was confirmed using FastQC version 0.11.8 (Andrews, 2010). To remove the “QuantSEQ FWD” adaptor sequence, the reads were trimmed using cutadapt version 1.18. The trimmed reads were mapped to the human reference genome (GRCh38.89) using the STAR aligning software v 2.6.0c. The RSEM software, version v1.3.1, was used to generate count tables. Genes were only retained if they were expressed at counts per million (cpm) above one in at least three samples. Counts were normalized using the TMM method (R-package edgeR, v3.36.0), followed by Voom transformation and differential expression analysis using Limma (R-package limma, v3.503.3). Principal component analysis showed that sample “CLBGA siPHF6a replicate 3” is an outlier so this sample was removed from further analysis. A general linear model was built with the treatment groups and the replicates as a batch effect. Gene Set Enrichment Analysis (GSEA) was performed on the genes ordered according to differential expression statistic value (t). Signature scores were conducted using a rank-scoring algorithm. GSEA was done using the Hallmark gene set collection from MSigDB (gsea-msigdb.org) or gene lists compiled from this or our previous study (GSE161902). All RNA-seq data are available through the Gene Expression Omnibus (GEO) repository (GSE228959). Signature scores were calculated based on the combined, ranked expressions levels of the genes in the geneset and then scored in the expression data.

For RT-qPCR, complementary DNA (cDNA) synthesis was done using the iScript Advanced cDNA synthesis kit from Bio-Rad. The PCR mix was composed out of 5 ng of cDNA, 2.5 µl of SsoAdvanced SYBR qPCR super mix (Bio-Rad), and 0.25 µl of forward and reverse primers (to a final concentration of 250 nM; Integrated DNA Technologies). The RT-qPCR cycling analysis was performed using a LC-480 device (Roche). qBasePlus software 3.2 (www.biogazelle.com) was used for the analysis of the gene expression levels. B2M, HPRT1, TBP, and HMBS were used as reference genes. The error bars in figures represent SD after error propagation, with mean centering and scaling to control. The primer designs are provided in **Table S5**.

### CUT&RUN

CUT&RUN coupled with high-throughput DNA sequencing was performed on isolated nuclei using Cutana pA/G-MNase (Epicypher, 15-1016) according to the manufacturer’s manual. Briefly, nuclei were isolated from 0.5M cells/sample in 100µl nuclear extraction buffer per sample and incubated with activated Concanavalin A beads for 10 min at 4°C while rotating. Nuclei were resuspended in 50µl antibody buffer containing a 1:100 dilution of each antibody (see **Table S3** for antibody information) and kept in an elevated angle on a nutator at 4°C overnight. Next, targeted digestion and release was performed with 2.5 µl Cutana pA/G-MNase (15-1116) and 100mM CaCl2 for 2 hours at 4°C on the nutator. After chromatin release by incubation on 37°C for 10 minutes, DNA was purified using the CUTANA DNA purification kit (14-0050) and eluted in 12µl of elution buffer. Sequencing libraries were prepared with the NEBNext Ultra II kit (Illumina, E7645), followed by paired-end sequencing on a Nextseq2000 using the NextSeq 2000 P2 Reagents 100 Cycles v3 (Illumina, 20046811). Prior to mapping on the human reference genome (GRCh37/hg19) with bowtie2 (v.2.3.1), quality of the raw sequencing data of CUT&RUN was evaluated using FastQC and adapter trimming was done using TrimGalore (v0.6.5). Quality of aligned reads were filtered using min MAPQ 30 and reads with known low sequencing confidence were removed using Encode Blacklist regions. For sample with a percentage duplicated reads higher then 10%, deduplication was performed using MarkDuplicates (Picard, v4.0.11). Peak calling was performed using MACS2 (v2.1.0) taking a q value of 0.05 as threshold and default parameters. Homer (v4.10.3) was used to perform motif enrichment analysis, with 200 bp around the peak summit as input. Overlap of peaks, annotation, heatmaps and pathway enrichment was analysed using DeepTools (v3.5.1), the R package ChIPpeakAnno (v3.28.1), and the web tool ENRICHR. The R package igvR (v1.19.3) was used for visualization of the data upon RPKM normalization. All CUT&RUN data are available through the Gene Expression Omnibus (GEO) repository (GSE228959). The detailed implementation is now available within in a Nextflow pipeline, accessible at https://github.com/PPOLLabGhent/nf_Pipeline_CUTandRUN.

## Data availability

- RNA seq data: Gene Expression Omnibus **GSE226288** (https://www.ncbi.nlm.nih.gov/geo/query/acc.cgi?acc=GSE228959)
- CUT&RUN seq data: Gene Expression Omnibus **GSE228958** (https://www.ncbi.nlm.nih.gov/geo/query/acc.cgi?acc=GSE228958)
- Modeling computer scrips: GitHub (https://github.com/PPOLLabGhent/nf_Pipeline_CUTandRUN)
- Protein Interaction IP-MS data: **PXD041025**

## Acknowledgements

We would like to thank prof. dr. Francis Impens and Sara Dufour for the technical assistance and subsequent data-analysis of the IP-MS experiments. This research was supported by the following funding agencies: Olivia Fund, ‘Kinderkankerfonds’, Ghent University (BOF.GOA.2022.0003.03 and BOF.STG.2022.0009.01) and Fund for Scientific Research Flanders (project G037918N). LD is supported by a UGent doctoral-assistant mandate (BOF22/PDO/024).

## Supplementary Figures

**Supplementary Figure S1.**
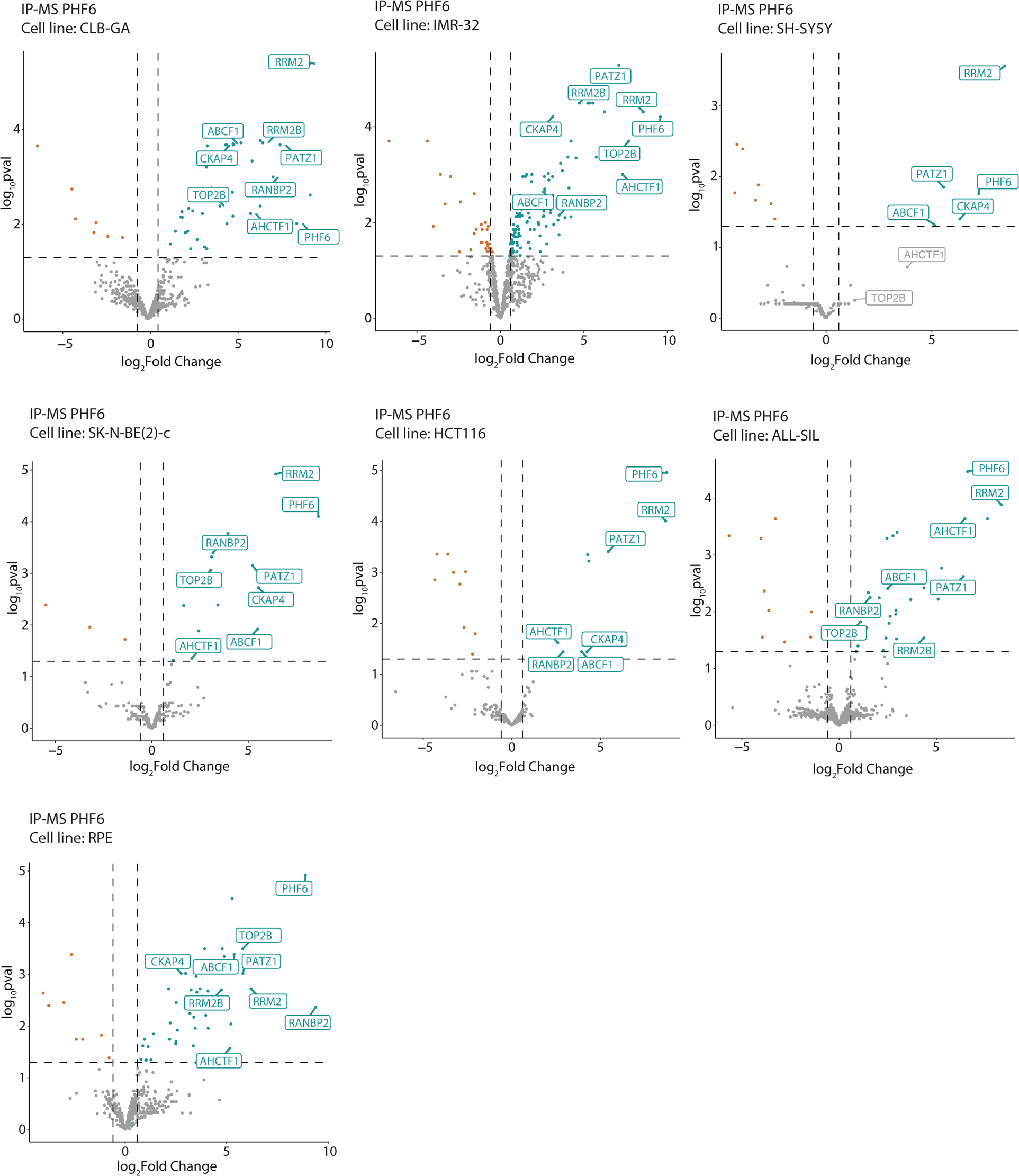
The PHF6-RRM2 interaction is robust pan-cancer as well as in non-malignant cell lines. Volcano plot of the immunoprecipitation coupled mass spectrometry (IP-MS) enriched hits in CLB-GA, IMR-32, SH-SY5Y, SK-N-BE(2)-C neuroblastoma cells, HCT116 colon cancer cells, ALL-SIL T-cell acute leukemia (T-ALL) cells and RPE, retinal epithelial cells using PHF6 as bait.

**Supplementary Figure S2.**
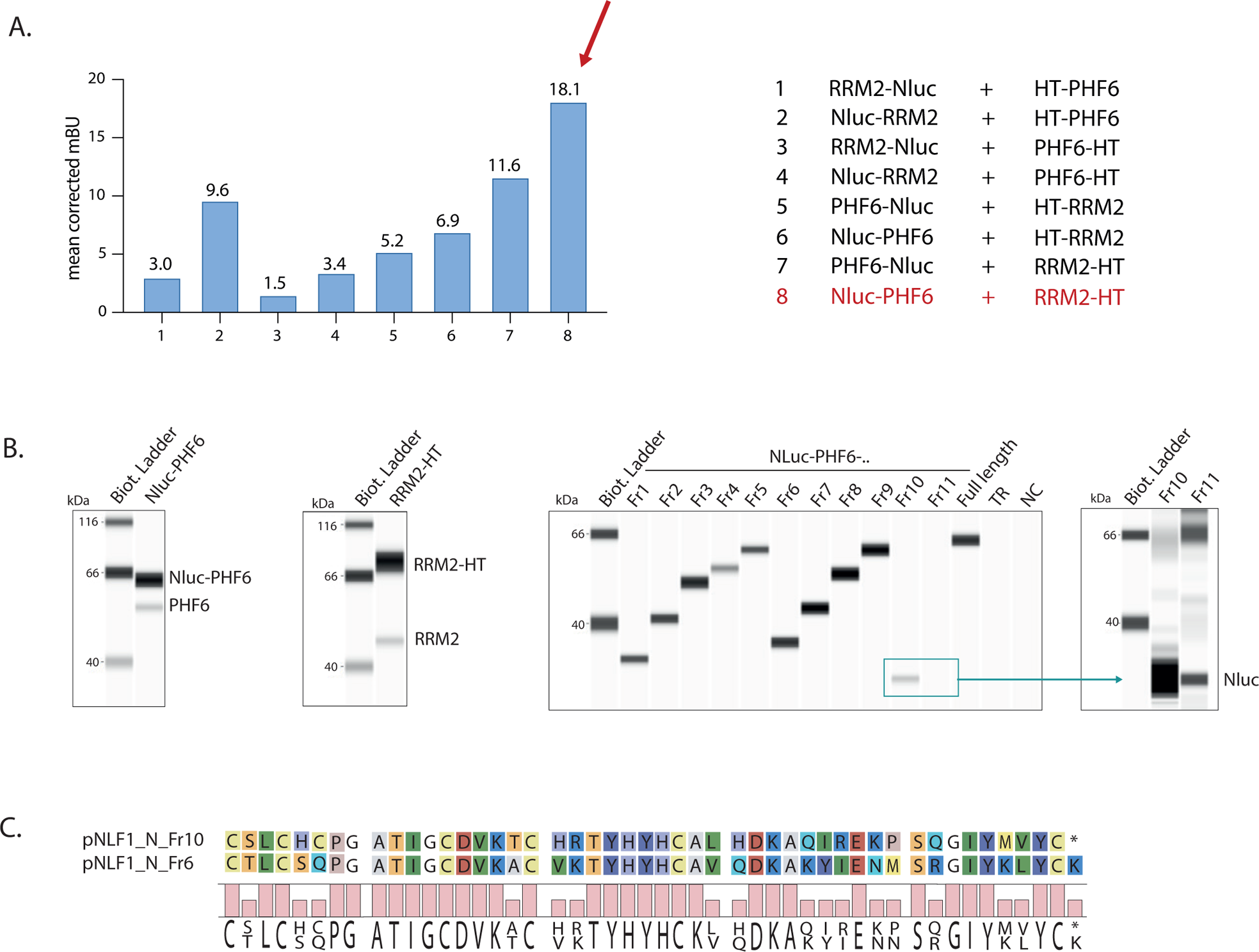
NanoBRET™ analysis confirms PHF6-RRM2 protein-protein interaction. **(A)** Mean corrected NanoBRET™ ratios resulting from transfecting different conformations of Nluc and HT fusion proteins in HEK293T cells. This revealed that the Nluc-PHF6 + RRM2-HT combination results in the highest NanoBRET™ ratio, which was selected for further experiments. **(B)** HEK293T cells were transfected with 2µg RRM2-HT, 0.2µg Nluc-PHF6 and 0.2µg of the deletion constructs Fr1-F11 fused to Nluc. Protein expression levels were analyzed with Simple WES using anti-RRM2, anti-PHF6 and anti-Nluc antibodies respectively. **(C)** Part of the protein sequence of the PHF6 deletion construct ‘fragment 10’ (Fr10, top) containing the first PHD domain and ‘fragment 6’ (Fr6, bottom) containing the second PHD domain of PHF6.

**Supplementary Figure S3.**
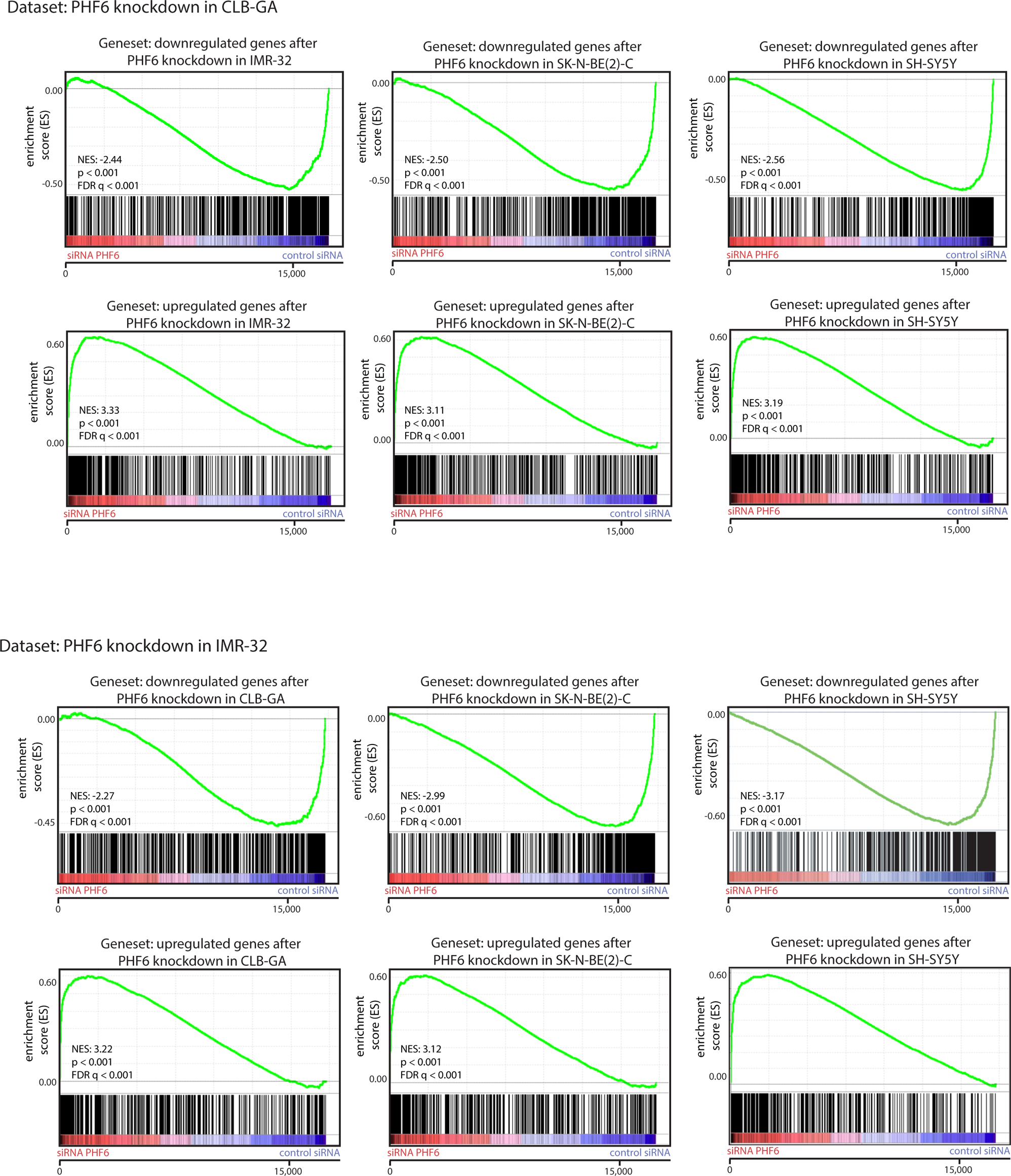
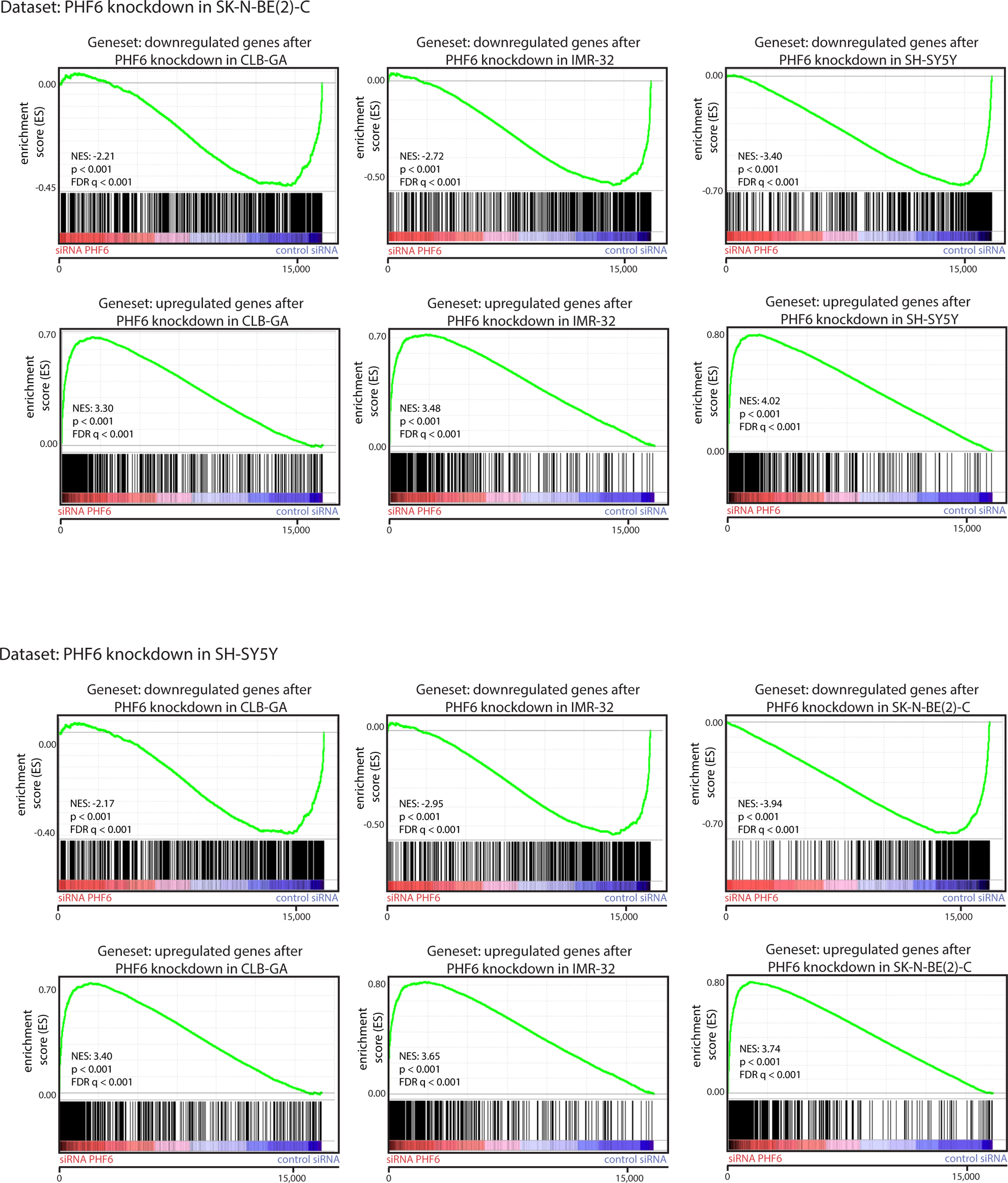
RNA-seq transcriptional profiles upon PHF6 knockdown in four neuroblastoma cell lines overlap significantly. GSEA shows a significant overlap between up- and down regulated genes upon PHF6 knockdown using siRNAs between all four neuroblastoma cell lines, CLB-GA, IMR-32, SK-N-BE(2)-C and SH-SY5Y.

**Supplementary Figure S4.**
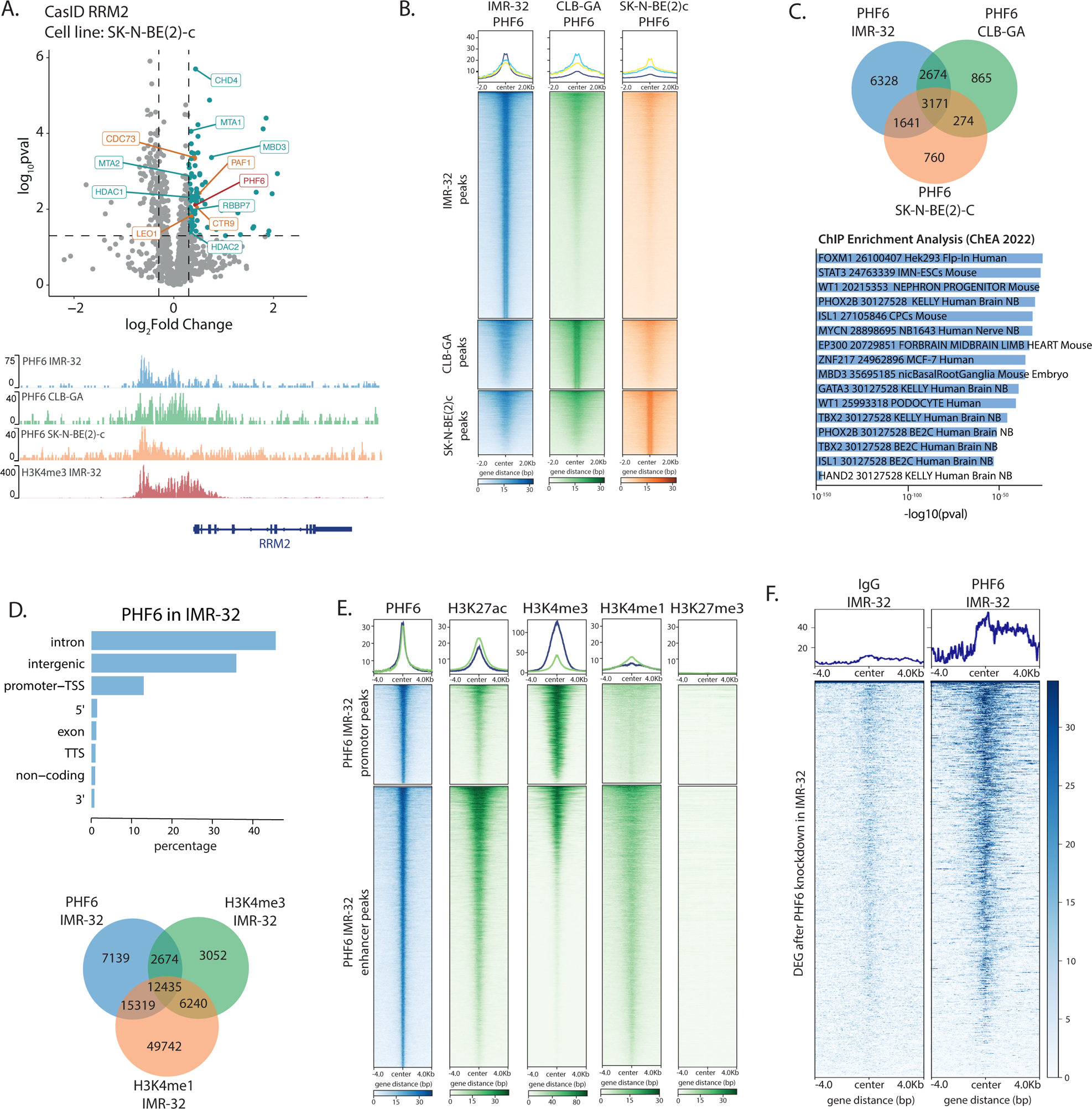
CUT&RUN analysis of PHF6 shows binding to active enhancers and promotors, including the RRM2 promotor. **(A)** Volcano plot of the Cas-ID experiment for the identification of RRM2 upstream regulators in SK-N-BE(2)-C cells (FDR < 0.05) (Nunes *et al*, 2022). PHF6 is depicted in red, PAF1 complex members in orange and NuRD complex members in blue (*top*), PHF6 and H3K3me3 binding activity at the *RRM2* locus in IMR-32, CLB-GA and SK-N-BE(2)-C cells from CUT&RUN experiments, confirming binding of PHF6 to the *RRM2* promotor region. Signals represent log likelihood ratio for the CUT&RUN signal compared to control signal (RPKM normalized). All peaks are called by MACS2 (q<0.05) (*bottom*); **(B)** Heatmap profiles −2kb and +2kb around the summit of PHF6 CUT&RUN peaks in IMR-32, CLB-GA and SK-N-BE(2)-C neuroblastoma cells, representing overlap of PHF6 binding sites in the three neuroblastoma cell lines. Peaks were ranked according to the sums of the peak scores across all datasets in the heatmaps; **(C)** Venn diagram showing the overlapping peaks between the three neuroblastoma cell lines (Fisher’s Exact Test p-value < 2.2 x 10^−16^) (*top*) and enrichment analysis using ENRICHR of the 3171 overlapping peaks in all cell lines reveals enrichment for ChIP-seq identified targets of amongst others FOXM1 and ADR CRC members ISL1, PHOX2B, HAND2, TBX2, GATA3 and MYCN (*bottom)*; **(D)** Peak annotation distribution analysis of PHF6 binding sites in IMR-32 cells (*top*) and Venn diagram showing overlap of PHF6 annotated peaks with peaks associated with the active promotor histone mark, H3K4me1 and active enhancer histone mark, H3K4me3 (Fisher’s Exact Test p-value < 2.2 x 10^−16^) (*bottom)*; **(E)** Heatmap profiles −4kb and +4kb around the summit of PHF6 CUT&RUN peaks, representing overlap of PHF6 with CUT&RUN peaks of H3K4me3, H3K27ac, H3K4me3 and H3K4me1 in IMR-32 neuroblastoma cells, grouped for promoters or enhancers (using homer annotation), and ranked according to the sums of the peak scores across all datasets in the heatmap; **(F)** Heatmap profiles −4kb and +4kb around the transcription start site of differentially expressed genes upon siRNA mediated PHF6 knockdown in IMR-32. On these regions, IgG control and PHF6 CUT&RUN data in IMR-32 are mapped and ranked according to the sums of the peak scores across all datasets in the heatmap.

## Supplementary Tables

**Supplementary Table 1.**
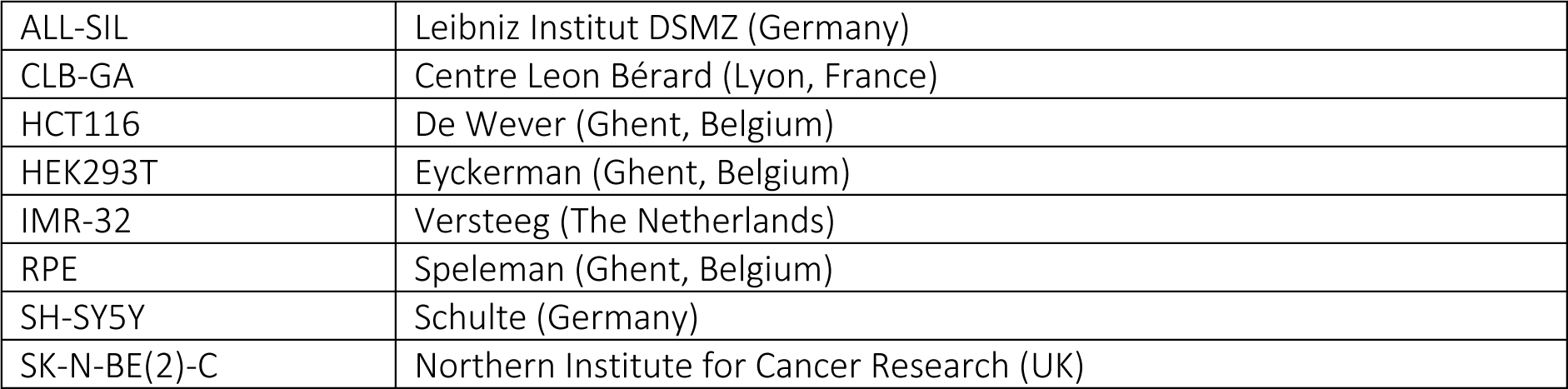
Origin of cell lines used in this manuscript.

|  |  |
| --- | --- |
| ALL-SIL | Leibniz Institut DSMZ (Germany) |
| CLB-GA | Centre Leon Bérard (Lyon, France) |
| HCT116 | De Wever (Ghent, Belgium) |
| HEK293T | Eyckerman (Ghent, Belgium) |
| IMR-32 | Versteeg (The Netherlands) |
| RPE | Speleman (Ghent, Belgium) |
| SH-SY5Y | Schulte (Germany) |
| SK-N-BE(2)-C | Northern Institute for Cancer Research (UK) |

**Supplementary Table 2.** Products used for cell culture and Western Blotting in this manuscript.

| Product | Obtained |
| --- | --- |
| 10x Tris/Glycine buffer | Bio-Rad Laboratories S.A.-N.V. |
| 10x Tris/Glycine/SDS buffer | Bio-Rad Laboratories S.A.-N.V. |
| 384- well plates | Corning |
| 96- well plates | Corning |
| b-mercapto-ethanol | Merck Life Science |
| BSA | Merck Life Science |
| Etoposide | Merck Life Science |
| Fetal bovine serum (FBS) | Merck Life Science |
| Immuno-Blot PVDF Membrane | Bio-Rad Laboratories S.A.-N.V. |
| L-Glutamine | Thermo Fisher Scientific BVBA |
| McCoy's 5A | LGC Standards SARL |
| Methanol | Thermo Fisher Scientific BVBA |
| Nitrocellulose membrane | Bio-Rad Laboratories S.A.-N.V. |
| Opti-MEM I Reduced Serum, no phenol red | Life Technologies Europe |
| PBS | Life Technologies Europe |
| Pen/Strep | Thermo Fisher Scientific BVBA |
| Phosphatase inhibitor mixture | Roche Molecular Biochemicals |
| Pierce™ BCA Protein Assay Kit | Life Technologies Europe |
| RPMI 1640 medium | Life Technologies Europe |
| SDS-PAGE gels, pre-cast | Bio-Rad Laboratories S.A.-N.V. |
| T25 flask | VWR |
| T75 flask | VWR |

**Supplementary Table 3.** Antibodies used in this manuscript.

| Protein | Company, antibody ID | Application & dilution |
| --- | --- | --- |
| ATR | Bethyl Laboratories, A300-138A | CUT&RUN, 1/100 |
| b-actin | Sigma-Aldrich, A2228 | Simple Wes, 1/150 |
| BRCA1 | Santa cruz, sc-646 | CUT&RUN, 1/100 |
| CHK1 | Santa cruz, sc-8408 | Western Blotting, 1/1000 |
| DDX11 | Abcam, ab230017 | CUT&RUN, 1/100 |
| FANCD2 | Biotechne, NB100-182 | CUT&RUN, 1/100 |
| H3K27ac | Thermo Fisher, MA5-24617 | CUT&RUN, 1/100 |
| H3K27me3 | Cell Signaling Technologies, #9733S | CUT&RUN, 1/100 |
| H3K4me1 | Abcam, ab8895 | CUT&RUN, 1/100 |
| H3K4me3 | Abcam, ab8580 | CUT&RUN, 1/100 |
| H3K56ac | Millipore, 07-677-I | coIP (ab1), 2µg |
| H3K56ac | Activ Motif, 39281 | coIP (ab2), 2µg<br>CUT&RUN, 1/100<br>Western Blotting, 1/1000 |
| gH2AX | Abcam, ab11174 | Western Blotting, 1/2000 |
| gH2AX | Bio-connect, GTX127342 | CUT&RUN, 1/100 |
| IgG | Abcam, ab2410 | IP-MS/coIP, 2µg |
| IgG | Cell Signaling Technologies, #3900 | CUT&RUN, 1/100 |
| MYCN | Santa Cruz, sc53993 | CUT&RUN, 1/100 |
| NanoLuc® | Promega, N7000 | Simple Wes, 1/50 |
| pCHK1 | Thermo Scientific, MA5-15145 | Western Blotting, 1/1000 |
| PHF6 | Bethyl Laboratories, A301-451A | IP-MS/coIP, 2µg<br>Western Blotting, 1/1000<br>CUT&RUN, 1/100 |
| PHF6 | Santa Cruz, sc365237 | Simple Wes, 1/100 |
| pRPA32/RPA2 (S33) | Abcam, ab211877 | Western Blotting, 1/500 |
| RPA32/RPA2 | Abcam, ab2175 | Western Blotting, 1/1000 |
| RRM2 | Abcam, ab154964 | Western Blotting, 1/1000 |
| RRM2 | Abcam, ab172476 | Simple Wes, 1/100 |
| Secondary Anti-Mouse Detection Module | Protein Simple, DM-002 | Simple Wes, undiluted according to manufactures protocol |
| Secondary Anti-Rabbit Detection Module | Protein Simple, DM-001 | Simple Wes, undiluted according to manufactures protocol |
| Secondary mouse HRP | Cell Signaling Technologies, #7076S | Western Blotting, 1/50 000<br>Simple Wes, 1/200 |
| Secondary rabbit HRP | Cell Signaling Technologies,, #7074S | Western Blotting, 1/50 000 |
| Veriblot rabbit secondary antibody | Abcam, ab131366 | Western Blotting, 1/5000 |
| Veriblot mouse secondary antibody | Abcam, ab131368 | Western Blotting, 1/5000 |
| Vinculin | Cell Signaling Technologies, #13901 | Western Blotting, 1/10000 |

**Supplementary Table 4.**
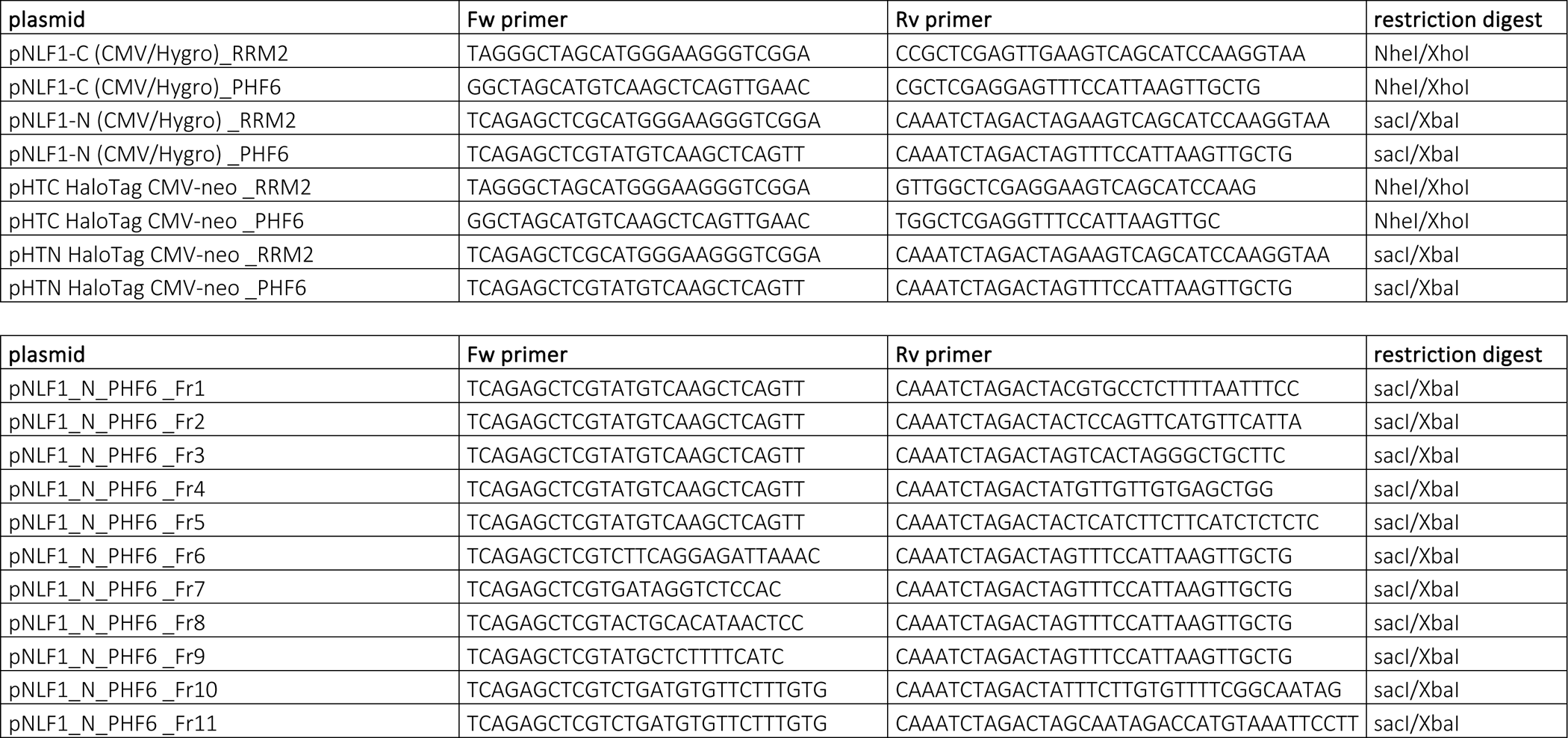
Primers and corresponding restriction digests used to clone the NanoBRET expression plasmids.

**Supplementary Table 5.**
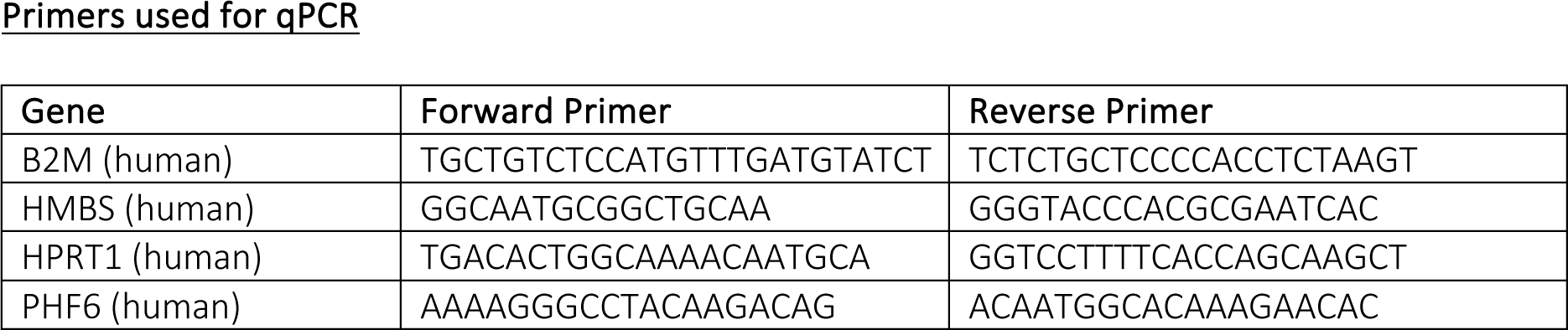
Sequences of primers used in this manuscript.

## References

Aasland R, Gibson TJ & Stewart AF (1995) The PHD finger: Implications for chromatin-mediated transcriptional regulation. Trends Biochem Sci 20: 56–59

Alvarez S, da Silva Almeida AC, Albero R, Biswas M, Barreto-Galvez A, Gunning TS, Shaikh A, Aparicio T, Wendorff AA, Piovan E, et al (2022) Functional mapping of PHF6 complexes in chromatin remodeling, replication dynamics, and DNA repair. Blood 139: 3418–3429

Battu A, Ray A & Wani AA (2011) ASF1A and ATM regulate H3K56-mediated cell-cycle checkpoint recovery in response to UV irradiation. Nucleic Acids Res 39: 7931–7945

Blosser WD, Dempsey JA, McNulty AM, Rao X, Ebert PJ, Lowery CD, Iversen PW, Webster YW, Donoho GP, Gong X, et al (2020) A pan-cancer transcriptome analysis identifies replication fork and innate immunity genes as modifiers of response to the CHK1 inhibitor prexasertib. Oncotarget 11: 216–236

Boeva V, Louis-Brennetot C, Peltier A, Durand S, Pierre-Eugène C, Raynal V, Etchevers HC, Thomas S, Lermine A, Daudigeos-Dubus E, et al (2017) Heterogeneity of neuroblastoma cell identity defined by transcriptional circuitries. Nat Genet 49: 1408–1413

Buisson R, Boisvert JL, Benes CH & Zou L (2015) Distinct but Concerted Roles of ATR, DNA-PK, and Chk1 in Countering Replication Stress during S Phase. Mol Cell 59: 1011–1024

Chao MM, Todd MA, Kontny U, Neas K, Sullivan MJ, Hunter AG, Picketts DJ & Kratz CP (2010) T-cell acute lymphoblastic leukemia in association with Börjeson-Forssman-Lehmann syndrome due to a mutation in PHF6. Pediatr Blood Cancer 55: 722–724

Che J, Smith S, Kim YJ, Shim EY, Myung K & Lee SE (2015) Hyper-Acetylation of Histone H3K56 Limits Break-Induced Replication by Inhibiting Extensive Repair Synthesis. PLoS Genet 11: 1–24

Choi EB, Vodnala M, Zerbato M, Wang J, Ho JJ, Inouye C, Ding L & Fong YW (2021) ATP-binding cassette protein ABCF1 couples transcription and genome surveillance in embryonic stem cells through low-complexity domain. Sci Adv 7

Cole KA, Huggins J, Laquaglia M, Hulderman CE, Russell MR, Bosse K, Diskin SJ, Attiyeh EF, Sennett R, Norris G, et al (2011) RNAi screen of the protein kinome identifies checkpoint kinase 1 (CHK1) as a therapeutic target in neuroblastoma. Proc Natl Acad Sci U S A 108: 3336–3341

Dale NC, Johnstone EKM, White CW & Pfleger KDG (2019) NanoBRET: The bright future of proximity-based assays. Front Bioeng Biotechnol 7: 56

Das C, Lucia MS, Hansen KC & Tyler JK (2009) CBP/p300-mediated acetylation of histone H3 on lysine 56. Nature 459: 113–117

Durbin AD, Zimmerman MW, Dharia N V., Abraham BJ, Iniguez AB, Weichert-Leahey N, He S, Krill-Burger JM, Root DE, Vazquez F, et al (2018) Selective gene dependencies in MYCN-amplified neuroblastoma include the core transcriptional regulatory circuitry. Nat Genet 50: 1240–1246

Feliciano YMS, Bartlebaugh JME, Liu Y, Rivera FJS, Bhutkar A, Weintraub AS, Buenrostro JD, Cheng CS, Regev A, Jacks TE, et al (2017) PHF6 regulates phenotypic plasticity through chromatin organization within lineage-specific genes. Genes Dev 31: 973–989

Fliedner A, Gregor A, Ferrazzi F, Ekici AB, Sticht H & Zweier C (2020) Loss of PHF6 leads to aberrant development of human neuron-like cells. Sci Rep 10: 1–15

Franz C, Walczak R, Yavuz S, Santarella R, Gentzel M, Askjaer P, Galy V, Hetzer M, Mattaj IW & Antonin W (2007) MEL-28/ELYS is required for the recruitment of nucleoporins to chromatin and postmitotic nuclear pore complex assembly. EMBO Rep 8: 165–172

Franzoni E, Booker SA, Parthasarathy S, Rehfeld F, Grosser S, Srivatsa S, Fuchs H, Tarabykin V, Vida I & Wulczyn FG (2015) miR-128 regulates neuronal migration, outgrowth and intrinsic excitability via the intellectual disability gene Phf6. Elife 4: 1–23

Gaggioli V, Lo CSY, Reverón-Gómez N, Jasencakova Z, Domenech H, Nguyen H, Sidoli S, Tvardovskiy A, Uruci S, Slotman JA, et al (2023) Dynamic de novo heterochromatin assembly and disassembly at replication forks ensures fork stability. Nat Cell Biol 25: 1017–1032

Van Groningen T, Koster J, Valentijn LJ, Zwijnenburg DA, Akogul N, Hasselt NE, Broekmans M, Haneveld F, Nowakowska NE, Bras J, et al (2017) Neuroblastoma is composed of two super-enhancer-associated differentiation states. Nat Genet 49: 1261–1266

Hajjari M, Salavaty A, Crea F & Kee Shin Y (2016) The potential role of PHF6 as an oncogene: a genotranscriptomic/proteomic meta-analysis. Tumor Biology 37: 5317–5325

Hay RT (2005) SUMO: A history of modification. Mol Cell 18: 1–12

Herold S, Kalb J, Büchel G, Ade CP, Baluapuri A, Xu J, Koster J, Solvie D, Carstensen A, Klotz C, et al (2019) Recruitment of BRCA1 limits MYCN-driven accumulation of stalled RNA polymerase. Nature 567: 545–549

Kamburov A, Wierling C, Lehrach H & Herwig R (2009) ConsensusPathDB—a database for integrating human functional interaction networks. Nucleic Acids Res 37: D623–D628

Kotsantis P, Petermann E & Boulton SJ (2018) Mechanisms of oncogene-induced replication stress: Jigsaw falling into place. Cancer Discov 8: 537–555 doi:10.1158/2159-8290.CD-17-1461 [PREPRINT]

Kuhn TM & Capelson M (2019) Nuclear pore proteins in regulation of chromatin state. Cells 8 doi:10.3390/cells8111414 [PREPRINT]

Kumar Jegadesan N & Branzei D DDX11 loss causes replication stress and pharmacologically exploitable DNA repair defects.

Li X, Yao H, Chen Z, Wang Q, Zhao Y & Chen S (2013) Somatic mutations of PHF6 in patients with chronic myeloid leukemia in blast crisis. Leuk Lymphoma 54: 671–672 doi:10.3109/10428194.2012.725203 [PREPRINT]

Liu Z, Li F, Ruan K, Zhang J, Mei Y, Wu J & Shi Y (2014) Structural and functional insights into the human börjeson-forssman-lehmann syndrome-associated protein PHF6. Journal of Biological Chemistry 289: 10069–10083

Loontiens S, Dolens AC, Strubbe S, Van de Walle I, Moore FE, Depestel L, Vanhauwaert S, Matthijssens F, Langenau DM, Speleman F, et al (2020) PHF6 Expression Levels Impact Human Hematopoietic Stem Cell Differentiation. Front Cell Dev Biol 8

Lower KM, Turner G, Kerr BA, Mathews KD, Shaw MA, Gedeon ÁK, Schelley S, Hoyme HE, White SM, Delatycki MB, et al (2002) Mutations in PHF6 are associated with Börjeson-Forssman-Lehmann syndrome. Nat Genet 32: 661–665

Mahtab M, Boavida A, Santos D & Pisani FM (2021) The genome stability maintenance DNA helicase DDX11 and its role in cancer. Genes (Basel) 12

Masumoto H, Hawke D, Kobayashi R & Verreault A (2005) A role for cell-cycle-regulated histone H3 lysine 56 acetylation in the DNA damage response. Nature 436: 294–298

Matsuoka S, Ballif BA, Smogorzewska A, McDonald ER, Hurov KE, Luo J, Bakalarski CE, Zhao Z, Solimini N, Lerenthal Y, et al (2007) ATM and ATR substrate analysis reveals extensive protein networks responsive to DNA damage. Science (1979) 316: 1160–1166

Mellor J (2006) It Takes a PHD to Read the Histone Code. Cell 126: 22–24

Murthy S & Reddy GPV (2006) Replitase: Complete machinery for DNA synthesis. J Cell Physiol 209: 711–717 doi:10.1002/jcp.20842 [PREPRINT]

Nunes C, Depestel L, Mus L, Keller KM, Delhaye L, Louwagie A, Rishfi M, Whale A, Kara N, Andrews SR, et al (2022) RRM2 enhances MYCN-driven neuroblastoma formation and acts as a synergistic target with CHK1 inhibition

Perez-Riverol Y, Bai J, Bandla C, García-Seisdedos D, Hewapathirana S, Kamatchinathan S, Kundu DJ, Prakash A, Frericks-Zipper A, Eisenacher M, et al (2022) The PRIDE database resources in 2022: A hub for mass spectrometry-based proteomics evidences. Nucleic Acids Res 50: D543–D552

Qiu S, Jiang G, Cao L & Huang J (2021) Replication Fork Reversal and Protection. Front Cell Dev Biol 9: 1–8

Rasmussen RD, Gajjar MK, Tuckova L, Jensen KE, Maya-Mendoza A, Holst CB, Møllgaard K, Rasmussen JS, Brennum J, Bartek J, et al (2016) BRCA1-regulated RRM2 expression protects glioblastoma cells from endogenous replication stress and promotes tumorigenicity. Nat Commun 7: N34

Ritchie ME, Phipson B, Wu D, Hu Y, Law CW, Shi W & Smyth GK (2015) limma powers differential expression analyses for RNA-sequencing and microarray studies. Nucleic Acids Res 43: e47–e47

Rother MB & van Attikum H (2017) DNA repair goes hip-hop: SMARCA and CHD chromatin remodellers join the break dance. Philosophical Transactions of the Royal Society B: Biological Sciences 372

Sarangi P & Zhao X (2015) SUMO-mediated regulation of DNA damage repair and responses. Trends Biochem Sci 40: 233

Saxena S & Zou L (2022) Hallmarks of DNA replication stress. Mol Cell 82: 2298–2314 doi:10.1016/j.molcel.2022.05.004 [PREPRINT]

Shendy NAM, Zimmerman MW, Abraham BJ & Durbin AD (2022) Intrinsic transcriptional heterogeneity in neuroblastoma guides mechanistic and therapeutic insights. Cell Rep Med 3: 100632 doi:10.1016/j.xcrm.2022.100632[PREPRINT]

Shimada M, Niida H, Zineldeen DH, Tagami H, Tanaka M, Saito H & Nakanishi M (2008) Chk1 Is a Histone H3 Threonine 11 Kinase that Regulates DNA Damage-Induced Transcriptional Repression. Cell 132: 221–232

Siegfried A, Rousseau A, Maurage CA, Pericart S, Nicaise Y, Escudie F, Grand D, Delrieu A, Gomez-Brouchet A, Le Guellec S, et al (2019) EWSR1-PATZ1 gene fusion may define a new glioneuronal tumor entity. Brain Pathology 29: 53–62

Técher H, Koundrioukoff S, Nicolas A & Debatisse M (2017) The impact of replication stress on replication dynamics and DNA damage in vertebrate cells. Nat Rev Genet 18: 535–550

Tian T, Bu M, Chen X, Ding L, Yang Y, Han J, Feng XH, Xu P, Liu T, Ying S, et al (2021) The ZATT-TOP2A-PICH Axis Drives Extensive Replication Fork Reversal to Promote Genome Stability. Mol Cell 81: 198–211.e6

Tjeertes JV., Miller KM & Jackson SP (2009) Screen for DNA-damage-responsive histone modifications identifies H3K9Ac and H3K56Ac in human cells. EMBO Journal 28: 1878–1889

Todd MAM, Ivanochko D & Picketts DJ (2015) Phf6 degrees of separation: The multifaceted roles of a chromatin adaptor protein. Genes (Basel) 6: 6

Todd MAM & Picketts DJ (2012) PHF6 interacts with the nucleosome remodeling and deacetylation (NuRD) complex. J Proteome Res 11: 4326–4337

Topal S, Vasseur P, Radman-Livaja M & Peterson CL (2019) Distinct transcriptional roles for Histone H3-K56 acetylation during the cell cycle in Yeast. Nat Commun 10

Van Vlierberghe P, Palomero T, Khiabanian H, Van Der Meulen J, Castillo M, Van Roy N, De Moerloose B, Philippé J, González-García S, Toribio ML, et al (2010) PHF6 mutations in T-cell acute lymphoblastic leukemia. Nat Genet 42: 338–342

Van Vlierberghe P, Patel J, Abdel-Wahab O, Lobry C, Hedvat C V, Balbin M, Nicolas C, Payer AR, Fernandez HF, Tallman MS, et al (2011) PHF6 mutations in adult acute myeloid leukemia. Leukemia 25: 130–134

Voss AK, Gamble R, Collin C, Shoubridge C, Corbett M, Gécz J & Thomas T (2007) Protein and gene expression analysis of Phf6, the gene mutated in the Börjeson-Forssman-Lehmann Syndrome of intellectual disability and obesity. Gene Expression Patterns 7: 858–871

Warmerdam DO, Alonso-de Vega I, Wiegant WW, van den Broek B, Rother MB, Wolthuis RM, Freire R, van Attikum H, Medema RH & Smits VA (2019) PHF6 promotes non-homologous end joining and G2 checkpoint recovery. EMBO Rep

Weiss WA, Aldape K, Mohapatra G, Feuerstein BG & Bishop JM (1997) Targeted expression of MYCN causes neuroblastoma in transgenic mice. EMBO Journal 16: 2985–2995

Whale AJ, King M, Hull RM, Krueger F & Houseley J (2022) Stimulation of adaptive gene amplification by origin firing under replication fork constraint. Nucleic Acids Res 50: 915–936

Whalen JM, Dhingra N, Wei L, Zhao X & Freudenreich CH (2020) Relocation of Collapsed Forks to the Nuclear Pore Complex Depends on Sumoylation of DNA Repair Proteins and Permits Rad51 Association. Cell Rep 31: 107635

Wieczorek D, Bögershausen N, Beleggia F, Steiner-Haldenstätt S, Pohl E, Li Y, Milz E, Martin M, Thiele H, Altmüller J, et al (2013) A comprehensive molecular study on coffin-siris and nicolaides-baraitser syndromes identifies a broad molecular and clinical spectrum converging on altered chromatin remodeling. Hum Mol Genet 22: 5121–5135

Williams SK, Truong D & Tyler JK (2008) Acetylation in the globular core of histone H3 on lysine-56 promotes chromatin disassembly during transcriptional activation. Proc Natl Acad Sci U S A 105: 9000–9005

De Wyn J, Zimmerman MW, Weichert-leahey N, Nunes C, Cheung BB, Abraham BJ, Beckers A, Volders PJ, Decaesteker B, Carter DR, et al (2021) Meis2 is an adrenergic core regulatory transcription factor involved in early initiation of th-mycn-driven neuroblastoma formation. Cancers (Basel) 13

Xiao W, Bharadwaj M, Levine M, Farnhoud N, Pastore F, Getta BM, Hultquist A, Famulare C, Medina JS, Patel MA, et al (2018) PHF6 and DNMT3A mutations are enriched in distinct subgroups of mixed phenotype acute leukemia with T-lineage differentiation. Blood Adv 2: 3526–3539

Xiao Z, Chang JG, Hendriks IA, Sigurdsson JO, Olsen J V. & Vertegaal ACO (2015) System-wide analysis of SUMOylation dynamics in response to replication stress reveals novel small ubiquitin-like modified target proteins and acceptor lysines relevant for genome stability. Molecular and Cellular Proteomics 14: 1419–1434

Xie Z, Bailey A, Kuleshov M V., Clarke DJB, Evangelista JE, Jenkins SL, Lachmann A, Wojciechowicz ML, Kropiwnicki E, Jagodnik KM, et al (2021) Gene Set Knowledge Discovery with Enrichr. Curr Protoc 1: e90

Yan Q, Wulfridge P, Doherty J, Fernandez-Luna JL, Real PJ, Tang HY & Sarma K (2022) Proximity labeling identifies a repertoire of site-specific R-loop modulators. Nat Commun 13

Yoo NJ, Kim YR & Lee SH (2012) Somatic mutation of PHF6 gene in T-cell acute lymphoblatic leukemia, acute myelogenous leukemia and hepatocellular carcinoma. Acta Oncol (Madr) 51: 107–111

Yuan J, Pu M, Zhang Z & Lou Z (2009) Histone H3-K56 acetylation is important for genomic stability in mammals. Cell Cycle

Zhang C, Mejia LA, Huang J, Valnegri P, Bennett EJ, Anckar J, Jahani-Asl A, Gallardo G, Ikeuchi Y, Yamada T, et al (2013) The X-Linked Intellectual Disability Protein PHF6 Associates with the PAF1 Complex and Regulates Neuronal Migration in the Mammalian Brain. Neuron 78: 986–993

Zhang X, Smits AH, Van Tilburg GBA, Ovaa H, Huber W & Vermeulen M (2018) Proteome-wide identification of ubiquitin interactions using UbIA-MS. Nature Protocols 2018 13:3 13: 530–550

